# Bycatch mitigation of endangered marine life

**DOI:** 10.1101/2024.02.20.581169

**Authors:** Mireia Villafáfila García, Antonio Carpio, Marga L Rivas

**Affiliations:** Department of Biology, Institute of Marine Science INMAR, University of Cádiz, Spain; Department of Zoology, Campus of Rabanales, University of Cordoba, 14071 Córdoba, Spain; Grupo de Sanidad y Biotecnología (SaBio), Instituto de Investigación en Recursos Cinegéticos, IREC (UCLM-CSIC-JCCM), Ronda Toledo 12, 13071 Ciudad Real, Spain

**Keywords:** Incidental fishing, cetaceans, LEDs, marine turtles, marine birds, rays, sea turtles, sharks

## Abstract

The fishing gear deployed by fishermen in seas and oceans throughout the world not only captures target species but also unintentionally ensnares non-target species, a phenomenon known as “by-catch”. This unintended capture of marine life can represent significant challenges for the fishing industry, with adverse impacts on both the environment and species such as sea turtles, marine mammals, seabirds and elasmobranchs, which may be injured or even killed. To address this problem, the fishing industry has implemented regulations and mitigation measures. In this literature review, we have examined 389 articles published between 2010 and 2022 that assess the effectiveness of these measures. It has been demonstrated that the most effective measures are ‘pingers’ for marine mammals, ‘TEDs’ (Turtle Excluder Devices) for sea turtles, and ‘BSLs’ (Bird Scaring Lines), more commonly known as ‘tori lines’, for seabirds. The most complex case is that of elasmobranchs, and the most effective measure has yet to be discovered. This complexity arises from the ongoing targeted fishing of these species, resulting in less monitoring of their catches and, therefore, fewer surveys. Overall, we encourage the global implementation of these measures by the fishing industry in order to reduce by-catch in an attempt to ensure the future of many endangered species.

## 1. INTRODUCTION

Fishing poses a significant threat to marine vertebrates, including sea turtles, marine mammals, seabirds and elasmobranchs. This threat stems from the impact of the fishing industry on these species, both as a direct target and incidentally. The underlying cause of this issue is the extensive presence of fishing fleets on the various seas and oceans. In 2020, the global fishing fleet was estimated to consist of approximately 4.1 million vessels, with Asia being the predominant continent, accounting for a total of 2.68 million vessels (FAO, 2022). These vessels utilize a range of fishing gear, depending on their target species. The most common gear types include gillnets, longlines, purse seines with and without purse lines, trawl nets and traps (FAO, 2022).

This fishing gear has a substantial impact on oceans and their ecosystems, resulting in a series of adverse effects. These include an overexploitation of species, such as overfishing, which can lead to the decline and even collapse of populations, ultimately affecting marine biodiversity and ecosystems (Crowder et al. 2008). Certain fishing gear also destroys habitats, as occurs with that used for bottom trawling since it damages coral reefs, seagrass beds and seabeds (Crowder et al. 2008). Furthermore, the disruption of the food web caused by the decline of specific species as a result of overfishing can affect other species dependent on them as a food source, leading to a cascading effect known as the ‘top-down’ or ‘bottom-up’ effect (Crowder et al. 2008). Additionally, non-target or by-catch species are often captured alongside the intended target species when non-selective fishing methods are employed, and a wide variety of species is captured, including those not intended for capture or trade (Crowder et al. 2008). This includes species that are commercially undesirable owing to their lower value or failure to comply with size or weight regulations, along with protected or endangered species (which may have serious implications for their conservation) and non-commercial species that are not of interest for trade (Agardy, 2000). In some cases, these non-target species are not only captured but also injured or even killed, and these species include marine mammals, sea turtles, seabirds and elasmobranchs (Crowder et al. 2008).

Certain management measures with which to mitigate the impacts on these species have been implemented, including catch and net restrictions and the deployment of specific fishing technologies tailored to the groups affected (Lucas and Berggren, 2023). In the case of sea turtles, physical and visual techniques have been tested, such as the Turtle Excluder Device (TED) (Warden, 2011) and Light Emitting Diodes (LEDs), which could also be a good alternative as regards mitigating sea turtle by-catch. In the case of marine mammals, acoustic, physical, visual and echolocation measures have also been studied, and acoustic deterrent devices such as pingers have been used. However, some studies have demonstrated that certain species may become habituated to pingers over time (Moan and Bjørge, 2021). With regard to seabirds, olfactory, physical and visual measures are used to mitigate by-catch, with Bird Scaring Lines (BSLs), more commonly known as tori lines, being the visual measure that has been most widely used in longline fisheries (Domingo et al. 2017). In the case of elasmobranchs, all types of measures have been tried, i.e., acoustic, olfactory, physical, visual, echolocation and electrosensory. However, the knowledge regarding their effectiveness is limited, as in the case of employing tori lines to reduce shark by-catch (Seidu et al. 2022; Jiménez et al. 2019).

In order to obtain a global overview, the principal objective of this study is to conduct a comprehensive review of the existing scientific literature focused on assessing the effectiveness of specific mitigation measures as regards reducing the by-catch of endangered species groups, including sea turtles, marine mammals, seabirds and elasmobranchs. As secondary objectives, this study aims to: (i) identify the mitigation measures most commonly employed to minimize by-catch on a global scale; (ii) analyze the geographic distribution of by-catch in fisheries by country in order to evaluate variations, and (iii) provide an overview of the worldwide mitigation measurements that have proven to have the greatest effectiveness as regards reducing by-catch for each group. The overall objective of this research is to contribute with valuable insights into mitigating the effect of by-catch on endangered marine species and to shed light on measures that have proven to be the most successful, along with areas in which further research is required.

## 2. MATERIALS AND METHODS

### 2.1. Literature review

This review was carried out using the Scopus, Web of Science and Google Scholar search engines to search for publications containing citation indices spanning from 2010 to 2022. The keywords used were a combination of “bycatch” AND “fisheries” AND “mitigation” AND “measures” AND “sea turtles” or “marine mammals” or “seabirds” or “elasmobranchs”. These four groups of marine vertebrates were selected as the objective of this study. The data collected were classified according to the animal class, species name in English, scientific name, UICN category, the mitigation measure used, the percentage of reduction in by-catch achieved, and finally, the reference of each paper (Table 1 SI).

**Table 1.**
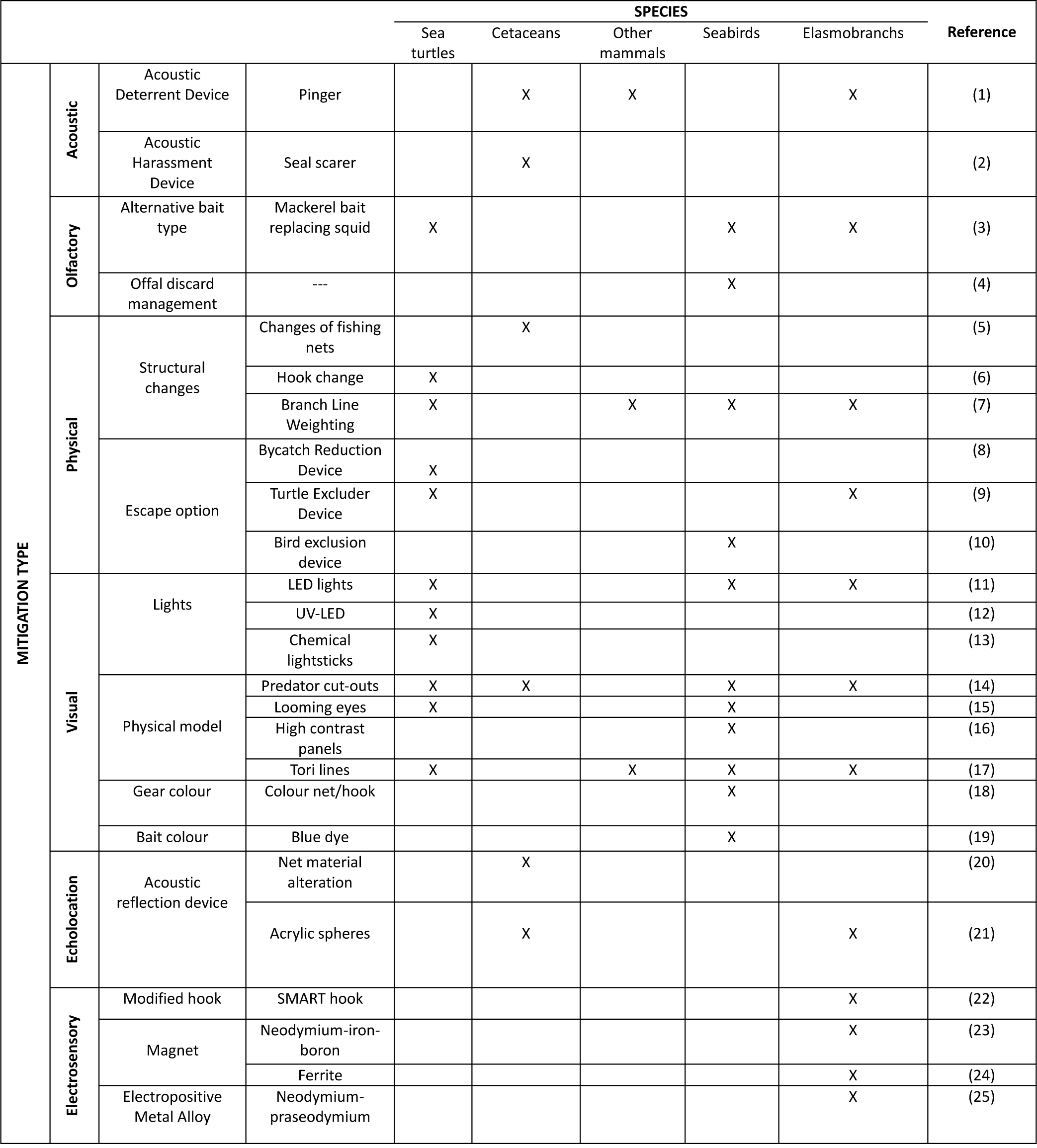
Mitigation measures proposed by different countries depending on the sensory system involved for each megafauna group, with their references (Methods SI).

|  |  |  |  | SPECIES |  |  |  |  | Reference |
| --- | --- | --- | --- | --- | --- | --- | --- | --- | --- |
|  |  |  |  | Sea turtles | Cetaceans | Other mammals | Seabirds | Elasmobranchs |  |
| MITIGATION TYPE | Acoustic | Acoustic Deterrent Device | Pinger |  | X | X |  | X | (1) |
|  |  | Acoustic Harassment Device | Seal scarer |  | X |  |  |  | (2) |
|  | Olfactory | Alternative bait type | Mackerel bait replacing squid | X |  |  | X | X | (3) |
|  |  | Offal discard management | --- |  |  |  | X |  | (4) |
|  | Physical | Structural changes | Changes of fishing nets |  | X |  |  |  | (5) |
|  |  |  | Hook change | X |  |  |  |  | (6) |
|  |  |  | Branch Line Weighting | X |  | X | X | X | (7) |
|  |  | Escape option | Bycatch Reduction Device | X |  |  |  |  | (8) |
|  |  |  | Turtle Excluder Device | X |  |  |  | X | (9) |
|  |  |  | Bird exclusion device |  |  |  | X |  | (10) |
|  | Visual | Lights | LED lights | X |  |  | X | X | (11) |
|  |  |  | UV-LED | X |  |  |  |  | (12) |
|  |  |  | Chemical lightsticks | X |  |  |  |  | (13) |
|  |  | Physical model | Predator cut-outs | X | X |  | X | X | (14) |
|  |  |  | Looming eyes | X |  |  | X |  | (15) |
|  |  |  | High contrast panels |  |  |  | X |  | (16) |
|  |  |  | Tori lines | X |  | X | X | X | (17) |
|  |  | Gear colour | Colour net/hook |  |  |  | X |  | (18) |
|  |  | Bait colour | Blue dye |  |  |  | X |  | (19) |
|  | Echolocation | Acoustic reflection device | Net material alteration |  | X |  |  |  | (20) |
|  |  |  | Acrylic spheres |  | X |  |  | X | (21) |
|  | Electrosensory | Modified hook | SMART hook |  |  |  |  | X | (22) |
|  |  | Magnet | Neodymium-iron-boron |  |  |  |  | X | (23) |
|  |  |  | Ferrite |  |  |  |  | X | (24) |
|  |  | Electropositive Metal Alloy | Neodymium-praseodymium |  |  |  |  | X | (25) |

### 2.2. Species and areas

We collected global data and information from scientific literature concerning the by-catch of the four marine vertebrate groups, such as sea turtles, marine mammals, seabirds and elasmobranchs, by fisheries in the different seas and oceans worldwide. Species of sea turtles included in the review belonged to the Cheloniidae and Dermochelyidae families. These species are distributed in seas and oceans throughout the world (except those in the Arctic and Antarctic), with the exception of Kemp’s ridley turtle (*Lepidochelys kempii*), which is solely a resident of the Gulf of Mexico and the northeastern coast of North America (Wibbels and Bevan, 2019).

Marine mammals such as odontocetes and mysticetes (families: Phocoenidae, Balaenopteridae, Delphinidae, Iniidae/Pontoporiidae), Sirenian and Pinniped (families: Dugongidae, Phocidae and Otariidae) were also included in the review. This group is, as a whole, widely distributed, as there are species that inhabit all the oceans in the world (including those in the Arctic and Antarctic), as in the case of the humpback whale (*Megaptera novaeangliae*) (Cooke, 2018). However, the majority of species are concentrated in the Atlantic, Pacific and Indian oceans, as is the case of the common dolphin (*Delphinus delphis*) (Braulik et al. 2021) and bottlenose dolphin (*Tursiops truncatus*) (Wells et al. 2019).

The seabird species included in our review belong to families Diomedeidae, Procellariidae, Spheniscidae, Anatidae, Phalacrocoracidae, Laridae, Sulidae, Stercorariidae and Oceanitidae. These birds are distributed across all continents, oceans and seas worldwide. Notably, the Mediterranean Sea serves as a prominent breeding and feeding area for several endangered species, such as the Balearic shearwater (*Puffinus mauretanicus*) (BirdLife International. 2018a) and the Yelkouan shearwater (*Puffinus yelkouan*) (BirdLife International. 2018b).

The review also encompassed elasmobranchs, which include sharks and rays from diverse families including Sphyrnidae, Triakidae, Carcharhinidae, Myliobatidae, Squalidae, Rajidae, Somniosidae, Rhinobatidae, Brachaeluridae, Lamnidae, Torpedinidae, Scyliorhinidae, Alopiidae, Pseudocarcharidae, Dasyatidae, Mobulidae, Glaucostegidae and Pentanchidae. Many of the species within this group are commonly found in coastal areas and inhabit the oceanic zone delimited by both tropics (Cancer and Capricorn). However, there are exceptions, such as the Greenland shark (*Somniosus microcephalus*), which inhabits the Arctic Circle (Kulka et al. 2020).

Overall, most of the sea turtle, marine mammal, seabird and elasmobranch species included in the study are classified by the UICN Red List of Threatened Species (UICN, 2023) as being Least concern (LC), Near threatened (NT), Vulnerable (VU), Endangered (EN), Critically Endangered (CR) and Data Deficient (DD) (Methods, Table 2 SI).

### 2.3. Statistical analysis

In order to verify the nature of the data obtained and compiled (Table 1 SI), the normality and homoscedasticity of the data were analyzed by employing R commander (version 2.7-0) using the Shapiro-Wilk test and Bartlett test, respectively. The data did not meet these criteria in any of the cases, and we therefore employed a non-parametric Kruskal-Wallis’s test. This test was used to examine whether there was a relationship (at the global level) between the percentage of reduction in by-catch and fishing areas, along with the percentage of reduction in by-catch per species group. In both cases, statistical significance was considered when p < 0.01. In order to carry out an individual analysis of the relationship between the efficiency of mitigation measures among the fisheries involved by group, a Goodness-of-Fit G-test was carried out by employing the R Studio program (version 4.0.2), using an R Script (Mangiafico, 2015) and the “DescTools” (Andri et al. 2021) and “RVAideMemoire” packages (Hervé, 2023).

## 3. RESULTS AND DISCUSSION

### 3.1. Overview of published literature

Of a total of 389 studies, 316 were excluded on the basis of specific criteria (Figure 1 SI). A total of 73 papers were eventually selected in order to analyze by-catch by year and fishery area. These are shown in Table 1 SI as follows: sea turtles in green, marine mammals in blue, seabirds in orange and elasmobranchs in purple. However, when attempting to assess the number of trials of fishing gear, it was possible to find only 31 studies that provided information on the number of trials (8140). Furthermore, studies related to proofs of concept or carried out in laboratories were not taken into account. The number of papers published each year that were found thanks to the literature search are shown in (Table 3 SI), with a maximum of 9 in 2018 and 2020. In contrast, only 3 studies were found in 2022 and 2 in 2010.

**Figure 1.**
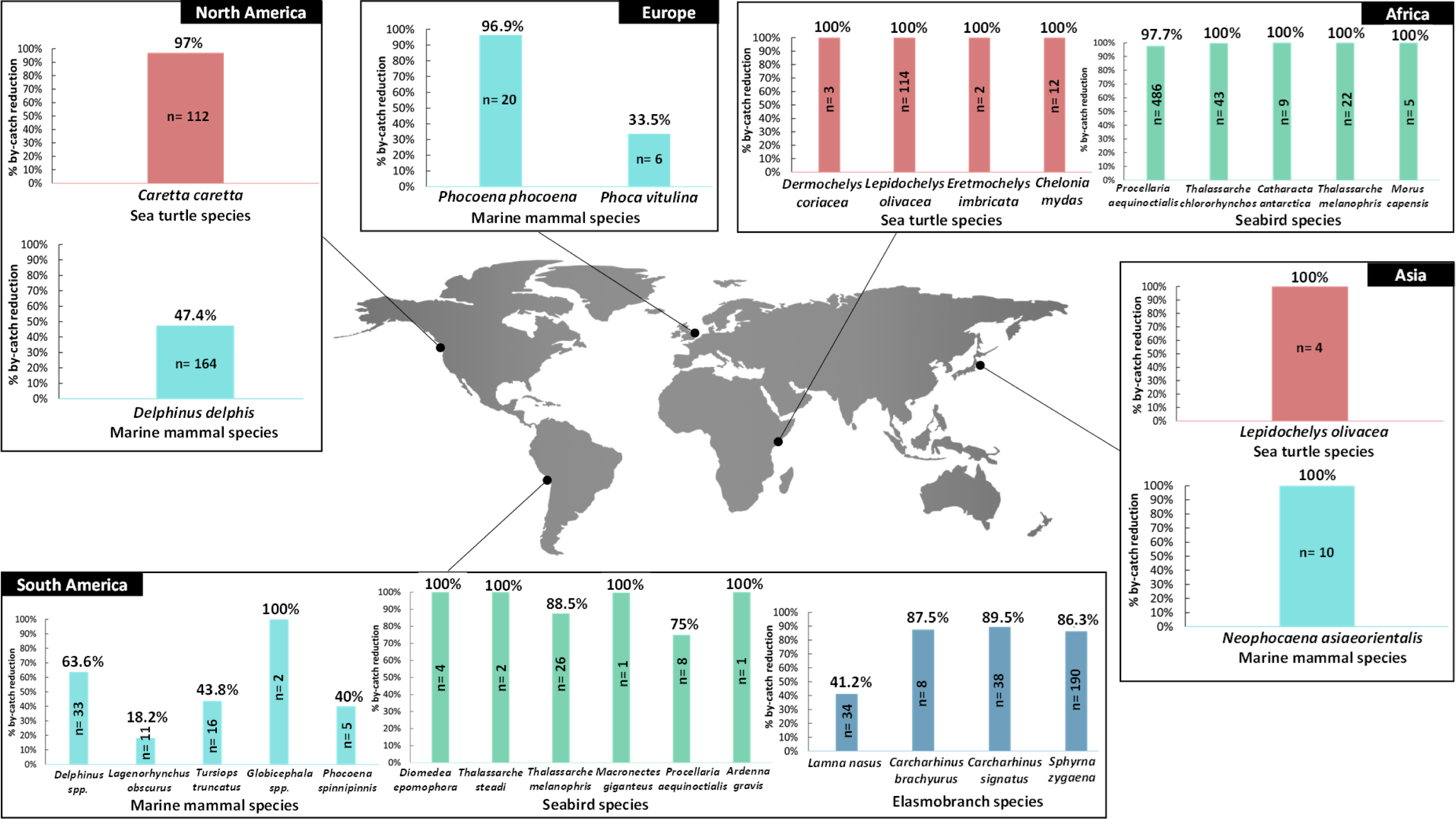
World map showing the percentage of reduction in by-catch per group (sea turtles in red, marine mammals in light blue, seabirds in green and elasmobranchs in dark blue) by continent (America divided into North (1) and South (2) America, Africa (3), Europe (4), Asia (5) and Oceania (6)), obtained by the most frequently used mitigation measure: TEDs for sea turtles, pingers for marine mammals and tori lines for seabirds and elasmobranchs.

The number of studies assessing by-catch by continent is shown in Figure 2 SI. The continent for which most studies had been conducted was America, with 28 studies, followed by Europe with 22. Within America, the United States carried out the largest number of studies, with 7 in total. In Europe, studies were conducted by 14 different countries, unlike that which occurred with Oceania, where all the studies were carried out in Australia. This figure also shows that there is a lack of studies in developing nations or locations in which small-scale fisheries (SSFs) are widespread, as is the case of Africa, where only 8 studies have been conducted. With regard to the number of trials carried out with each type of fishing gear, gillnets dominated the count with a total of 7636 trials, distantly followed by set net with 273 trials (Figure 3 SI).

**Figure 2.**
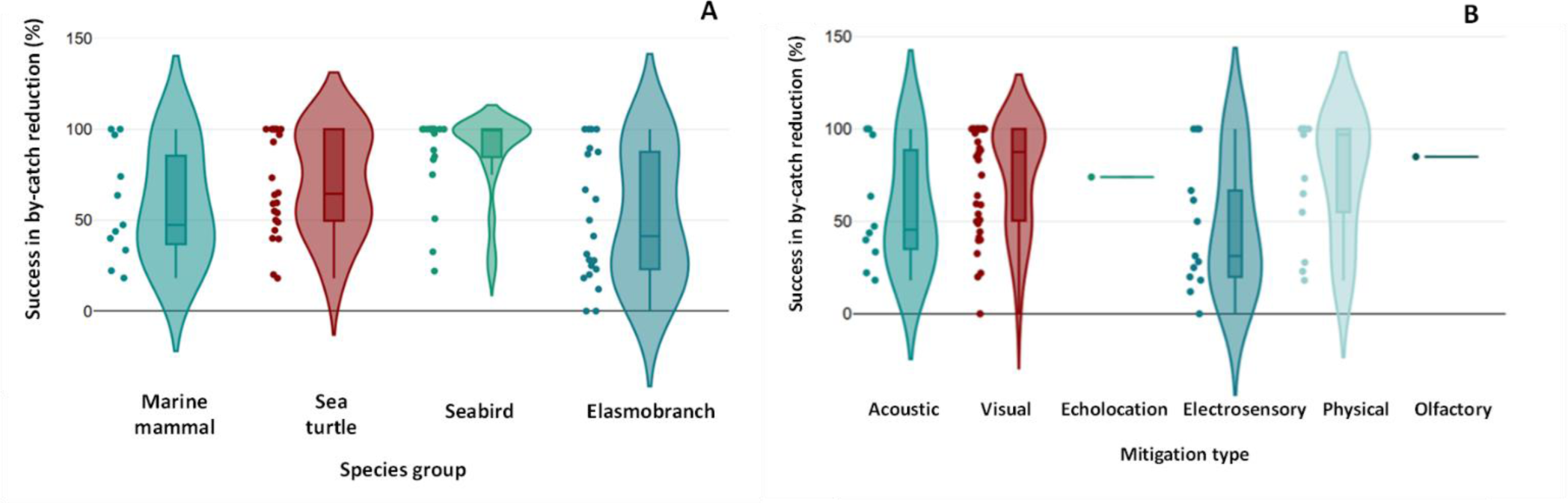
A. Violin plot illustrating the percentage of reduction in by-catch obtained for the species groups (marine turtles, marine mammals, seabirds and elasmobranchs). B. Percentage of reduction in by-catch obtained with the different mitigation types (acoustic, visual, echolocation, electrosensory, physical and olfactory). The horizontal lines inside each box correspond to the mean, while the vertical lines at the ends of each box refer to standard deviation (SD).

**Figure 3.**
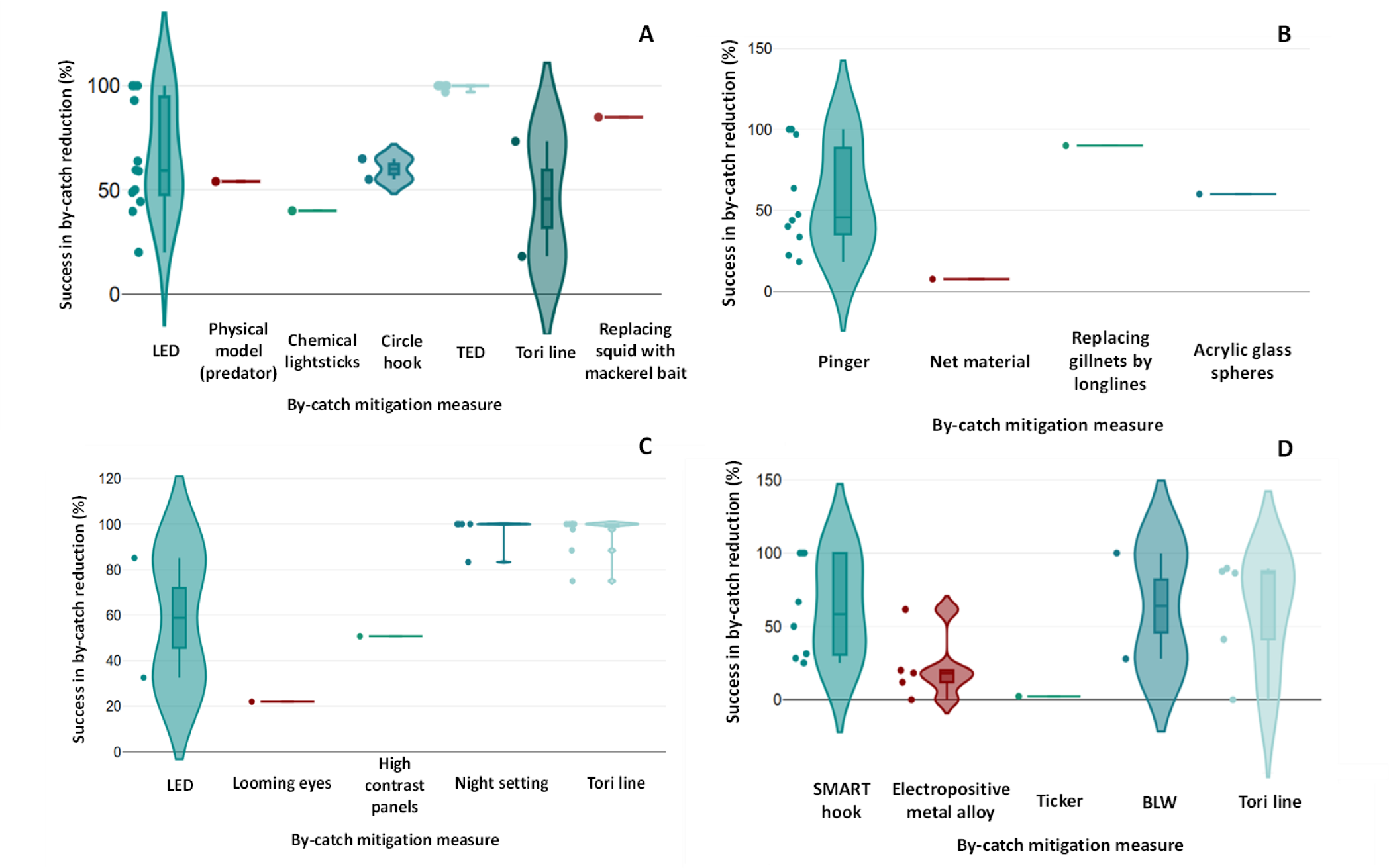
Success in reducing by-catch (%) of **A.** Sea turtles; **B.** Marine mammals; **C.** Seabirds, and **D**. Elasmobranchs, using different by-catch mitigation measures. The horizontal lines inside each box correspond to the mean, while the vertical lines at the ends of each box refer to SD.

### 3.2. Worldwide mitigation measures

Several mitigation measures with which to minimize the by-catch of these megafauna species have been tested in different fisheries around the world. A summary of the mitigation measures, categorized by the sensory system involved for each group, is provided in Table 1. In the case of sea turtles, the most commonly used olfactory measure involved using an alternative bait type. With regard to visual measures, LED lights emerged as a promising alternative, while in the case of physical measures, various escape options such as TEDs were frequently employed. In the case of odoncetes, mysticetes and other marine mammals such as the harbour seal (*Phoca vitulina*) and the Californian sea lion (*Zalophus californianus),* the measures most frequently used were pingers, a type of Acoustic Deterrent Devices (ADDs). With regard to seabirds, the mitigation measure that predominated was the tori line, a type of seabird scaring device (Gilman et al. 2021) consisting of a line towed from a high point at the aft of a vessel, from which several streamers are attached to scare seabirds and prevent their access to the critical area where baited hooks sink (Domingo et al. 2017). The main mitigation measure used for elasmobranchs was electrosensory devices. However, there was no single predominant measure, with SMART hooks (Grant et al. 2018; O’Connell et al. 2014), rare earth (Porsmoguer et al. 2015; Westlake et al. 2018), ferrite magnets (Richards et al. 2018) and Electropositive Metal (EPM) Alloy (Godin et al. 2013; Hutchinson et al. 2012) all being employed. However, it seems that tori lines have obtained good results for some of the species in this group (Jiménez, Forselledo, et al. 2019).

### 3.3. Mitigation measures by group

Overall, it was found that, globally, there were significant differences among the percentage of reduction in by-catch per fishery area (χ=36.33, df=19, p<0.01, n=20, Kruskal-Wallis’s test) (Figure 1) and in the percentage of success in by-catch per group (χ=14.67, df=3, p<0.01, n=4, Kruskal-Wallis’s test) (Figure 2A).

However, no significant differences were found as regards the percentage of reduction in by-catch resulting from different mitigation types (χ=10.17, df=4, p=0.04, n=6, Kruskal-Wallis’s test) (Figure 2B).

#### 3.3.1. Sea turtles

In the case of sea turtles, there was no significant difference among the percentage of reduction in by-catch according to the fishery area (G=9.15, X-squared df=10, p=0.52, n=11, G-test). Notably, the US Mid-Atlantic bottom trawl gear (Warden, 2011) and the industrial trawling fishery of Gabon (Casale et al. 2017) attained the highest values of reduction in by-catch per individual (mean ± SD, 0.97 ± NA and 1.0 ± NA, respectively). Similarly, no significant differences were observed among sea turtle species (G=7.46, X-squared df=4, p=0.11, n=24, G-test). However, the species with the highest percentages of reduction in by-catch reduction were the loggerhead turtle (*Caretta caretta*) (n=112) and the olive ridley turtle (*Lepidochelys olivacea*) (n=114) (97% and 100%, respectively). This was achieved using physical measures, specifically TEDs (Figure 1).

##### 3.3.1.1. Olfactory

One olfactory measure was investigated by testing the use of alternative bait types to mitigate incidental sea turtle by-catch in longline fisheries. The best results were attained by replacing squid bait with mackerel bait (Figure 3A), reducing by-catch by 88% and 85% for all turtles species (Coelho et al. 2012). In addition, by combining the effects of the mackerel by exchanging the traditional J-style hooks for two circle hooks (one non-offset and one with 10° offset), it was possible to obtain a 50-59% reduction in the by-catch (n=223) (Coelho et al. 2015).

##### 3.3.1.2. Physical

Tori lines were used in a Uruguayan pelagic longline fishery to reduce the by-catch. A reduction of 18.1% was obtained for the loggerhead turtle (n=83), while one of 73.3% was obtained for the leatherback turtle (*Dermochelys coriacea*) (n=15) (Jiménez et al. 2019) (Figure 3A).

TEDs have also been tested in numerous trawl fisheries, as occurred in the US Mid-Atlantic, with a reduction in by-catch of 97% for the loggerhead turtle (n=112) (Warden, 2011). A reduction in by-catch of 100% was documented for Indian fisheries (Raghu et al. 2016), and a similar reduction of 100% was achieved for four sea turtle species (n=131) in Gabonese fisheries (Casale et al. 2017) (Figure 1).

##### 3.3.1.3. Visual

Of all the possible visual mitigations methods for sea turtles, those that have been studied most are LED lights. However, there are also others, such as chemical lightsticks, physical models (predator cut-outs) and buoyless nets (Figure 3A).

The implementation of 500 nm green LEDs in an Indonesian small-scale coastal gillnet fishery led to a reduction in multi-species sea turtle by-catch of 61.4% (n=10), and specifically 59.5% of green turtle (*Chelonia mydas*) (n=14) (Gautama et al. 2022). This same measure was also used in a Mexican gillnet fishery, obtaining a reduction of 50% for the loggerhead turtle (n=17) (Senko et al., 2022) and 59% (n=85), 63.9% (n=125) and 48.8% (n=41) for green turtles (Wang et al. 2010; Ortiz et al. 2016; Kakai, 2019). Chemical lightsticks, meanwhile, obtained a reduction of 40% for the green turtle (n=85) (Wang et al. 2010). The implementation of 100-400 nm LED lights in a small-scale gillnet fishery led to a decrease in green turtle by-catch of 93% (n=13) (Darquea et al. 2020). In Italian and Mexican fisheries, the use of LEDs led to a reduction in loggerhead turtle by-catch of 100% (n=18) (Lucchetti et al. 2019; Virgili et al. 2018), and a reduction in green turtle by-catch of 39.7% (n=209) when using UV net illumination (Wang et al. 2013). Although TEDs have attained the best results, more trials on LEDs should be included, as they seem to be a good alternative by which to reduce the by-catch of these species.

#### 3.3.2. Marine mammals

In the case of marine mammals, there were no significant differences among the percentages of reduction in by-catch according to fishery areas (G=5.00, X-squared df=5, p=0.41, n=6, G-test). Notably, the highest values of by-catch reduction per individual were attained by the small set net Japanese fishery (Amano et al. 2017) and Norwegian commercial fisheries (Moan and Bjørge, 2021) (mean ± SD, 10.0 ± NA and 13.0 ± 9.8, respectively). Similarly, no significant differences were found among marine mammal species (G=0.57, X-squared df=9, p=0.99, n=11, G-test). However, the narrow-ridged finless porpoise (*Neophocaena asiaeorientalis*) (n=10) and harbour porpoise (*Phocoena phocoena*) (n=20) were the species with the highest percentage of reduction in by-catch (100% and 96.9%, respectively) when using acoustic measures, specifically pingers (Figure 1).

##### 3.3.2.1. Acoustic

Sensory technologies, specifically acoustic reflectors and pingers, were designed in the late 1970s and 1980s to deter marine mammals in gillnet fisheries (Dawson, 1991).

The AQUAmark 100 pinger, which operates at between 20 and 160kHz, achieved a reduction in narrow-ridged finless porpoise by-catch of 100% (n=10) (Amano et al. 2017) (Figure 1). The long-term effectiveness of the Dukane Netmark 1000 Pinger, which operates at 12-100 kHz harmonics, was assessed in a gillnet fishery (Carretta and Barlow, 2011). However, only the by-catch of two species decreased, specifically the common dolphin by 47.4% and the Northern elephant seal (*Mirounga angustirostris*) by 80.8% (n=164). When used in a driftnet fishery, the same pinger achieved reductions of between 18.2% and 100%, depending on the species (Mangel et al. 2013).

The evaluation of two pingers in gillnet fisheries, i.e., the Banana pinger by Fishtek Marine Industries (operating at 50-120 kHz, 154 dB) and the Dolphin pinger by Future Oceans (operating at 70 kHz, 132 dB), attained positive results, with reductions of 96.90% and 33.50% (Moan and Bjørge, 2021) (Figure 1). Furthermore, the implementation of these devices did not have a major negative impact on their daily fishing operations and contributed to the reduction in marine mammal by-catch.

##### 3.3.2.2. Physical

Berninsone et al. (2020) replaced gillnets with longlines in order to minimize the by-catch of franciscana (*Pontoporia blainvillei*), which it is considered the most threatened cetacean in the South Western Atlantic (Negri et al. 2012; Bordino and Albareda, 2004). This study decreased the by-catch of this species by 90% (n=85) (Figure 3B), thus showing that this method is an excellent alternative. However, only one study was carried out, which makes it difficult to generalize its effectiveness to other areas.

##### 3.3.2.3. Visual

Unlike that which occurs with sea turtles, visual mitigation measures are not commonly used to prevent marine mammal by-catch. Tori lines and Bird Line Weighting (BLW) have been tested in Uruguayan pelagic longline fisheries with no significant results (Jiménez et al. 2019). In order to evaluate the real effect of these measures on marine mammals, further research is consequently necessary.

##### 3.3.2.4. Echolocation reflection

Mysticetes and odontocetes use echolocation, which helps them to determine the location of objects in the sea. A mitigation measure consisting of adding acrylic glass spheres to a gillnet has consequently been developed in order to reduce the by-catch of the harbour porpoise (n=5), obtaining a reduction of 60% (Kratzer et al. 2021). Another measure tested was the modification of two types of nets, a barium sulfate net and a stiff nylon net (Bordino et al. 2013), but a reduction of only 7.4% (n=54) was obtained (Figure 3B).

#### 3.3.3. Seabirds

With regard to seabirds, the percentage of reduction in by-catch was not significantly different among fishery areas (G=7.84, X-squared df=5, p=0.16, n=6, G-test). Similarly, no significant differences were found among seabird species (G=1.58, X-squared df=16, p=1, n=20, G-test). Notably, the species with the highest percentage of by-catch reduction were the Atlantic yellow-nosed albatross (*Thalassarche chlororhynchos*) (n=43), the black-browed albatross (*Thalassarche melanophris*) (n=22) and the white-chinned petrel (*Procellaria aequinoctialis*) (n=486) (100%, 100% and 97.7%, respectively) when using visual measures, and specifically tori lines (Figure 1).

##### 3.3.3.1. Olfactory

The main olfactory mitigation measures used to minimize seabird by-catch were offal discard management (Kuepfer et al. 2022; Collins et al. 2021; Rollinson et al. 2017), thawed bait (Collins et al. 2021; Rollinson et al. 2017), blue-dyed bait (Gilman et al. 2021), artificial bait (Cortés and González-Solís, 2018) and replacing squid with mackerel as bait (Gonzalez et al. 2012; Li et al. 2012) (**¡Error! No se encuentra el origen de la referencia.**). These measures were, in certain instances, reinforced with non-sensory methods, such as night setting (Collins et al. 2021; Rollinson et al. 2017), seasonal closures (Collins et al. 2021), hook management (Collins et al. 2021) and the limitation of by-catch rates per year (Rollinson et al. 2017).

Despite the variety of existing measures, there is limited information regarding their effectiveness as regards reducing seabird by-catch and their effect on commercial catches. For instance, the evaluation of artificial bait demonstrated a reduction in target catches of 77% when compared to control lines (Cortés and González-Solís, 2018), but sample sizes were not included in the study.

##### 3.3.3.2. Physical

In order to reduce the by-catch rate of seabirds by using physical measures, the increase in the sink rate of baited hooks by reducing the distance between the hook and the weight of the branch lines (65g) was tested in a pelagic longline fishery (Jiménez et al. 2019), obtaining a reduction of 42.5%. Others studies propose the introduction of BLW as a mitigation measure, such as that by Paterson et al. (2019), which was carried out in a demersal longline fishery where a reduction in the by-catch was from 90.9% to 100%.

##### 3.3.3.3. Visual

The measures most commonly used in the case of seabirds are those of visual mitigation and include techniques such as LEDs (Bielli et al. 2020; Mangel et al. 2018; Field et al. 2019), high contrast panels (Field et al. 2019; Oliveira et al. 2021), buoys with looming eyes (Rouxel et al. 2021), night setting (Cortés and González-Solís, 2018) and tori lines (Cortés and González-Solís, 2018; Gilman et al. 2021) (**¡Error! No se encuentra el origen de la referencia.**).

The implementation of 500 nm LEDs was positive as regards reducing the by-catch of 4 species (n=46), with a reduction of 84% (Bielli et al. 2020), while green LEDs led to a reduction of 85.1% (Mangel et al. 2018). However, in the study carried out by Field et al. (2019), the efficacy of two types of 500 nm LEDs (constant green lights and flashing white LED lights) achieved a reduction of only 32.6% (n=43), although the use of high contrast panels reduced the by-catch of species by 50.8% (n=65). New devices such as the “Looming eyes buoy” (LEB) have also emerged, leading to a decrease in the by-catch of seabirds species of 22% (n=5724) (Rouxel et al. 2021) (Figure 3C).

Another measure is that of night setting, which has been tested with the artisanal demersal longliners of the Western Mediterranean. Although the sample sizes were limited, the results obtained showed a reduction of 83.3% and 100% (n=19) (Figure 3C), and an increase in sample testing is, therefore, recommended in order to ensure the efficiency of this measure.

Finally, the most widely used measure with which to reduce the by-catch of seabirds is that of tori lines, which have been shown to provide a significant reduction in the by-catch of all birds species, from 97.7% to 100% (Domingo et al. 2017; Paterson et al. 2019) (Figure 1).

#### 3.3.4. Elasmobranchs

With regard to elasmobranchs, the percentage of reduction in by-catches was significantly different according to the study site (G=11.83, X-squared df=2, p<0.01, n=3, G-test), with a Uruguayan longline fishery attaining the highest values as regards a reduction in by-catch (Jiménez et al. 2019). No significant differences were found among elasmobranch species (G= 1.60, X-squared df=17, p=1, n=21, G-test), although the species that attained the highest percentage of reduction in by-catch were the night shark (*Carcharhinus signatus)* (n=38) and the smooth hammerhead (*Sphyrna zygaena*) (n=190) (89.5% and 86.3%, respectively) when using visual measures such as tori lines (Figure 1).

##### 3.3.4.1. Acoustic

During the study of the effectiveness of pingers at reducing the by-catch of certain species of marine mammals, their effect was also analyzed for elasmobranchs. No significant differences in captures were attained when using Aquamark 100 and 200 pingers (Bilgin and Kose, 2018; Mangel et al. 2013).

##### 3.3.4.2. Olfactory

Only one olfactory mitigation measure with which to reduce the by-catch of different species of sharks and rays has been studied over a 13-year period (Coelho et al. 2012): that of replacing squid with mackerel bait. However, the effectiveness of this measure was limited.

##### 3.3.4.3. Physical

Physical measures by which to mitigate the by-catch of elasmobranchs include BLW (Jiménez et al. 2019), Bycatch Reduction Devices (BRDs) (Gupta et al. 2020) and the use of a ‘tickler’ (Kynoch et al. 2015), i.e., a piece of chain placed in front of the bottom gear of the trawler that is considered effective as regards catching skates and rays that may escape under the net. The inclusion of BLW in a Uruguayan longline fishery led to a reduction in the by-catch of the scalloped hammerhead (*Sphyrna lewini*) (n=2) of 100%, while the figure for the pelagic stingray (*Pteroplatytrygon violacea*) (n=18) was 27.8% (Figure 3D). However, in the case of the “tickler”, the number of species captured increased for all species with the exception of the lesser-spotted dogfish (*Scylorhinus canicular*) (n=1525), which attained a decrease of 2.3% (Kynoch et al. 2015) (Figure 3D).

##### 3.3.4.4. Visual

The use of LED lights as a mitigation measure has been tested for all four groups (sea turtles, marine mammals, seabirds and elasmobranchs), with an uncertain effect on elasmobranchs (Mangel et al. 2018). However, in a Mexican gillnet fishery, there was a reduction in the elasmobranch by-catch of 95% (Senko et al. 2022). Tori lines have also obtained good results for this group, reducing the by-catch rate for the porbeagle (*Lamna nasus*) (n=34), copper shark (*Carcharhinus brachyurus*) (n = 8), night shark (n=38) and smooth hammerhead (n=190) by 41.2%, 87.5%, 89.5% and 86.3%, respectively (Jiménez et al. 2019) (Figure 1).

##### 3.3.4.5. Echolocation

Rays do not use echolocation, and the by-catch data obtained after the implementation of the acrylic glass spheres by Kratzer et al. (2021) in a Turkish commercial fishery confirm this. More thornback skate (*Raja clavata*) individuals were caught in the modified gillnet (n=97) than in the standard one (n=41).

##### 3.3.4.6. Electrosensory

Sharks have a complex and extensive electrosensory system, which includes the ampullae of Lorenzini located around the snout or rostral area (Kajiura and Holland, 2002). The use of SMART hooks in a longline fishery in the Gulf of Maine (USA) led to a reduction in the number of shark species caught, from 25% to 100% (O’Connell et al. 2014) (Figure 3D).

The last sensory type measure found was the use of hooks made from a neodymium-praseodymium alloy, use by longlines in USA and Ecuador with the scalloped hammerhead (n=52), leading to a reduction of 61.5% (Hutchinson et al. 2012) (Figure 3D).

### 3.4. Limitations of this review

The objective of this literature review was to provide a comprehensive overview of the most effective measures used to date in order to reduce the by-catch of sea turtles, marine mammals, seabirds and elasmobranchs. However, it was difficult to carry out the global standardization of data because many studies were incomplete owing to a lack of sample sizes (Königson et al. 2022; Kuepfer et al. 2022; Godin et al. 2013), the existence of small sizes (O’Connell et al. 2014; Domingo et al. 2017; Jiménez et al. 2019) or the absence of the name of the species being studied (*Diomedea spp., Procellaria spp., Delphinus spp., Globicephala spp.*) (Yokota et al. 2011; Mangel et al. 2013). Moreover, in many cases the effectiveness of different mitigation measures for the species involved was not included (Kynoch et al. 2015; Porsmoguer et al. 2015; Basran et al. 2020), making it difficult to make comparisons among studies.

### 3.5. Recommendations for future research

A limited number of complete studies on by-catch mitigation measures were found. These included TEDs for sea turtles (Warden, 2011; Casale et al. 2017), pingers for marine mammals (Amano et al. 2017; Moan and Bjørge, 2021) and tori lines for seabirds (Domingo et al. 2017; Paterson et al. 2019). However, the available research on this topic is still lacking in many aspects.

There is a need for more studies that quantitatively assess the actual amount of by-catch (Basran et al. 2020; Culik et al. 2015; Westlake et al. 2018). Moreover, most of the results obtained often vary according to geographical areas, species and fishing practices, thus highlighting the importance of conducting further research into effective strategies by which to mitigate by-catch, particularly in regions in which SFFs are prevalent, such as Asia, Africa and South America.

It is also essential to establish standardized reporting practices, define study parameters, specify research locations and context, and examine unintended impacts on animal populations so as to attain accurate comparisons (Kynoch et al. 2015; Porsmoguer et al. 2015; Grant et al. 2018). Consistency in measurement metrics is crucial, focusing on the number of individuals captured per unit of effort (Senko et al. 2022; Berninsone et al. 2020; Gautama et al. 2022). Thorough documentation should encompass specifics such as gear type, study locale, the technology employed, and technical specifications (Senko et al. 2022; Mangel et al. 2013). It is, therefore, also recommended that studies explicitly detail sample and effect sizes (O’Connell et al. 2014; Domingo et al. 2017).

Furthermore, we believe that it is necessary to increase exploration into the combination of sensory deterrents in order to reduce by-catch across various taxonomic groups (Coelho et al. 2015; Gilman et al. 2021). Future research should prioritize the use of cost-efficient technologies that are straightforward to implement, as these are more likely to gain the support and compliance of the fishing industry. This will make it possible to work toward preserving the future of many endangered species and reducing the impact of by-catch.

## Author Contributions

Conceptualization, A.J.C., M.L.R.: data cleansing and writing— original draft preparation, M.V.; writing—review and supervision, A.J.C., M.L.R. All authors have read and agreed to the published version of the manuscript.

## Acknowledgments

M.V.G. is supported by the ‘Plan propio de estímulo y apoyo a la Investigación y Transferencia. INICIA-INV: Iniciación a la investigación (2022-2023)’ of the University of Cadiz. A.J.C. is supported by a ‘Juan de la Cierva’ contract (IJC2020-042629-I) funded by MCIN/AEI/10.13039/501100011033 and by the European Union Next GenerationEU/PRTR. M.L.R. was supported by Postdoctoral Research Contracts of the Andalusian government (Spain) and FEDER EU funds. This work received financial support from the Agencia Andaluza de Cooperación Internacional para el Desarrollo AACID (projects EMEP 2022UI007_2022).

