## Supplementary material for "Bycatch mitigation of endangered marine life": Supplementary_information.pdf

### 2. Material and methods

### 2.1. Literature review

**Table 1 SI.** Compilation of relevant information (class, species, scientific name, UICN category, % of reduction in by-catch, mitigation measure and reference) from the 73 papers studied in this review. Sea turtles appear in green, marine mammals in blue, seabirds in orange and elasmobranchs in purple.

| | Class | Species | Scientific name | UICN category | % of by-catch reduction | Mitigation measure | Reference |
|---|---|---|---|---|---|---|---|
| 1 | Marine mammal | Narrow-ridged finless porpoise | *Neophocaena asiaeorientalis* | Endangered | 100% | Pinger | Amano, M., Kusumoto, M., Abe, M., Akamatsu, T. (2017). Long-term effectiveness of pingers on a small population of finless porpoises in Japan. Endangered Species Research, 32, 35-40. https://doi.org/10.3354/esr00776 |
| 2 | Marine mammal | Humpback whale | *Megaptera novaeangliae* | Least Concern | NA | Pinger/seal scarer | Basran, CJ., Woelfng, B., Neumann, C., Rasmussen, MH. (2020). Behavioural responses of Humpback Whales (*Megaptera novaeangliae*) to two acoustic deterrent devices in a Northern feeding ground of Iceland. Aquat Mamm 46(6):584–602. https://doi.org/10.1578/AM.46.6.2020.584 |
| 3 | Marine mammal | Duskydolphin | *Lagenorhynchus obscurus* | Least Concern | 70.8% | LED lights | Bielli, A., Alfaro-Shigueto, J., Doherty, PD., Godley, BJ., Ortiz, C., Pasara, A., Wang, JH., Mangel, JC. (2020). An illuminating idea to reduce bycatch in the Peruvian small-scale gillnet fishery. Biol Cons. https://doi.org/10.1016/j.biocon.2019.108277 |
| | | Burmeister's porpoise | *Phocoena spinipinnis* | Near Threatened | | | |
| | Sea turtle | Leatherback turtle | *Dermochelys coriacea* | Vulnerable | 74.4% | | |
| | | Hawksbill turtle | *Eretmochelys imbricata* | Critically Endangered | | | |
| | | Loggerhead turtle | *Caretta caretta* | Vulnerable | | | |
| | | Green turtle | *Chelonia mydas* | Endangered | | | |
| | | Olive ridley turtle | *Lepidochelys olivacea* | Vulnerable | | | |
| | Seabird | Waved albatross | *Phoebastria irrorata* | Critically Endangered | 84% | | |
| | | Pink-footed shearwater | *Ardenna creatopus* | Vulnerable | | | |
| | | White-chinned petrel | *Procellaria aequinoctialis* | Vulnerable | | | |
| | | Humboldt penguin | *Spheniscus humboldti* | Vulnerable | | | |

| | | | | | | | |
|---|---|---|---|---|---|---|---|
| | Elasmobranch | Smooth hammerhead | *Sphyrna zygaena* | Vulnerable | NA | | |
| | | NA | *Mustelus spp.* | NA | | | |
| | | Copper shark | *Carcharhinus brachyurus* | Vulnerable | | | |
| | | Blue shark | *Prionace glauca* | Near Threatened | | | |
| | | NA | *Myliobatis spp.* | NA | | | |
| 4 | Marine mammal | Harbour porpoise | *Phocoena phocoena* | Least Concern | 0% | Pinger | Bilgin, S., Kose, O (2018). Testing two types of acoustic deterrent devices (pingers) to reduce harbour porpoise, *Phocoena phocoena* (Cetacea: Phocoenidae) by catch in turbot (Psetta maxima) set gillnet fishery in the Black Sea, Turkey. Cahiers De Biol Marine 59(5): 473–479. https://doi.org/10.21411/CBM.A.D5B58D5B |
| | Elasmobranch | Spiny dogfish | *Squalus acanthias* | Vulnerable | 0% | | |
| | | Thornback skate | *Raja clavata* | Near Threatened | 0% | | |
| 5 | Marine mammal | Harbour porpoise | *Phocoena phocoena* | Least Concerned | NA | Pinger | Königson, S., Naddaf, R., Hedgärde, M., Pettersson, A., Östman, Ö., Benavente, E., Amundin, M. (2021). Will harbor porpoises (*Phocoena phocoena)* be deterred by a pinger that cannot be used as a "dinner bell" by seals? Marine Mamm Sci. https://doi.org/10.1111/mms.12880 |
| 6 | Marine mammal | Harbour porpoise | *Phocoena phocoena* | Least Concern | 60% | Acrylic glass spheres | Kratzer, I, Brooks, M., Bilgin, S., Ozdemir, S., Kindt-Larsen, L., Larsen, F., Stepputtis, D (2021). Using acoustically visible gillnets to reduce bycatch of a small cetacean: first pilot trials in a commercial fishery. Fish Res. https://doi.org/10.1016/j.fshres.2021.106088 |
| | Elasmobranch | Thornback skate | *Raja clavata* | Near Threatened | 0% | | |
| 7 | Sea turtle | Green turtle | *Chelonia mydas* | Endangered | 59.5% | LED lights | Gautama, D. A., Susanto, H., Riyanto, M., Wahju, R. I., Osmond, M., Wang, J. H. (2022). Reducing sea turtle bycatch with net illumination in an Indonesian small-scale coastal gillnet fishery. Frontiers in Marine Science, 9, 1036158. https://doi.org/10.3389/fmars.2022.1036158 |
| | | Hawksbill turtle | *Eretmochelys imbricata* | Critically Endangered | 61.4% | | |
| | | Olive ridley turtle | *Lepidochelys olivacea* | Vulnerable | | | |
| | | Leatherback turtle | *Dermochelys coriacea* | Vulnerable | | | |
| | | Loggerhead turtle | *Caretta caretta* | Vulnerable | | | |
| | | Flatback turtle | *Natator depressus* | Data Deficient | | | |
| 8 | Seabird | Black-browed albatross | *Thalassarche melanophris* | Least Concern | NA | Offal discard | Kuepfer, A., Sherley, R. B., Brickle, P., Arkhipkin, A., Votier, S. C. (2022). Strategic discarding reduces seabird numbers and contact rates with trawl fishery gears in the Southwest Atlantic. Biological Conservation, 266, 109462. https://doi.org/10.1016/j.biocon.2022.109462 |
| | | NA | *Macronectes spp.* | NA | NA | | |
| 9 | Elasmobranch | NA | NA | NA | 95% | LED lights | Senko, JF., Peckham, SH., Aguilar-Ramirez, D., Wang, JH. (2022). Net illumination reduces fisheries bycatch, maintains catch value, and increases operational eficiency. Curr Biol 32:1–8. https://doi.org/10.1016/j.cub.2021.12.050 |
| | Sea turtle | Loggerhead turtle | *Caretta caretta* | Vulnerable | 50% | | |
| 10 | Marine mammal | Harbour porpoise | *Phocoena phocoena* | Least Concern | 79.7% | Pinger | Chladek, J., Culik, B., Kindt-Larsen, L., Albertsen, CM., von Dorrien, C (2020). Synthetic harbour porpoise (*Phocoena phocoena*) communication signals emitted by acoustic alerting device (Porpoise ALert, PAL) significantly reduce their bycatch in western Baltic gillnet |

| | | | | | | | |
|---|---|---|---|---|---|---|---|
| | | | | | | | fisheries. Fish Res. https://doi.org/10.1016/j.fshres.2020.105732 |
| 11 | Sea turtle | Loggerhead turtle | *Caretta caretta* | Vulnerable | 100% | UV-LED | Lucchetti, A., Bargione, G., Petetta, A., Vasapollo, C., Virgili, M. (2019). Reducing sea turtle bycatch in the Mediterranean mixed demersal fisheries. Frontiers in Marine Science, 6, 387. https://doi.org/10.3389/fmars.2019.00387 |
| | | Loggerhead turtle | *Caretta caretta* | Vulnerable | NA | TED | |
| 12 | Marine mammal | Harbour porpoise | *Phocoena phocoena* | Least Concern | 9% | Pinger | Omeyer, L., Doherty, P. D., Dolman, S., Enever, R., Reese, A., Tregenza, N., ... Godley, B. J. (2020). Assessing the effects of banana pingers as a bycatch mitigation device for harbour porpoises (*Phocoena phocoena*). Frontiers in Marine Science, 285. https://doi.org/10.3389/fmars.2020.00285 |
| 13 | Seabird | Long-tailed duck | *Clangula hyemalis* | Vulnerable | 22% | Looming eyes | Rouxel, Y., Crawford, R., Cleasby, I. R., Kibel, P., Owen, E., Volke, V., ... Oppel, S. (2021). Buoys with looming eyes deter seaducks and could potentially reduce seabird bycatch in gillnets. Royal Society open science, 8(5), 210225. https://doi.org/10.1098/rsos.210225 |
| 14 | Seabird | Long-tailed duck | *Clangula hyemalis* | Vulnerable | 32.6% | LED | Field, R., Crawford, R., Enever, R., Linkowski, T., Martin, G., Morkūnas, J., ... Oppel, S. (2019). High contrast panels and lights do not reduce bird bycatch in Baltic Sea gillnet fisheries. Global Ecology and Conservation, 18, e00602.https://doi.org/10.1016/j.gecco.2019.e00602 |
| | | Velvet scoter | *Melanitta fusca* | Vulnerable | 50.8% | High contrast panels | |
| 15 | Sea turtle | Loggerhead turtle | *Caretta caretta* | Vulnerable | 100% | UV-LED | Virgili, M., Vasapollo, C., Lucchetti, A. (2018). Can ultraviolet illumination reduce sea turtle bycatch in Mediterranean set net fisheries? Fisheries Research, 199, 1-7. https://doi.org/10.1016/j.fishres.2017.11.012 |
| 16 | Marine mammal | Franciscana | *Pontoporia blainvillei* | Vulnerable | 90% | Changing fishing nets | Berninsone, L. G., Bordino, P., Gnecco, M., Foutel, M., Mackay, A. I., Werner, T. B. (2020). Switching gillnets to longlines: an alternative to mitigate the bycatch of Franciscana Dolphins (*Pontoporia blainvillei*) in Argentina. Frontiers in Marine Science, 7, 699. https://doi.org/10.3389/fmars.2020.00699 |
| 17 | Sea turtle | Olive ridley turtle | *Lepidochelys olivacea* | Vulnerable | No significative | LED lights | Darquea, J. J., Ortiz-Alvarez, C., Córdova-Zavaleta, F., Medina, R., Bielli, A., Alfaro-Shigueto, J., Mangel, J. C. (2020). Trialing net illumination as a bycatch mitigation measure for sea turtles in a small-scale gillnet fishery in Ecuador. Latin american journal of aquatic research, 48(3), 446-455. http://dx.doi.org/10.3856/vol48-issue3-fulltext-2428 |
| | | Green turtle | *Chelonia mydas* | Endangered | 93% | | |
| | | Leatherback turtle | *Dermochelys coriacea* | Vulnerable | NA | | |
| 18 | Marine mammal | Humpback whale | *Megaptera novaeangliae* | Least Concern | NA | Pinger | Erbe, C., McPherson, C. (2012). Acoustic characterisation of bycatch mitigation pingers on shark control nets in Queensland, Australia. Endangered Species Research, 19(2), 109-121. https://doi.org/10.3354/esr00467 |
| | | NA | *Dolphin spp* | NA | NA | | |
| | | Dugong | *Dugong dugon* | Vulnerable | NA | | |
| 19 | Marine mammal | Franciscana | *Pontoporia blainvillei* | Vulnerable | 7.4% | Net material (Barium sulfate or stiff nylon) | Bordino, P., Mackay, A. I., Werner, T. B., Northridge, S. P., Read, A. J. (2013). Franciscana bycatch is not reduced by acoustically reflective or physically stiffened gillnets. Endangered Species Research, 21(1), 1-12. https://doi.org/10.3354/esr00503 |
| | | Common dolphin | *Delphinus delphis* | Least Concern | 47.4% | | |
| | | Northern right whale dolphin | *Lissodelphis borealis* | Least Concern | No significative | | |

| | | | | | | | |
|---|---|---|---|---|---|---|---|
| 20 | Marine mammal | Pacific white-sided dolphin | *Lagenorhynchus obliquidens* | Least Concern | No significative | Pinger | Carretta, J. V., Barlow, J. (2011). Long-term effectiveness, failure rates, and “dinner bell” properties of acoustic pingers in a gillnet fishery. Marine Technology Society Journal, 45(5), 7-19. https://doi.org/10.4031/MTSJ.45.5.3 |
| | | Northern elephant seal | *Mirounga angustirostris* | Least Concern | 80.8% | | |
| | | Californian sea lion | *Zalophus californianus* | Least Concern | 0% | | |
| 21 | Elasmobranch | Greenland shark | *Somniosus microcephalus* | Vulnerable | NA | SMART hook | Grant, S. M., Sullivan, R., Hedges, K. J. (2018). Greenland shark (*Somniosus microcephalus*) feeding behavior on static fishing gear, effect of SMART (Selective Magnetic and Repellent-Treated) hook deterrent technology, and factors influencing entanglement in bottom longlines. PeerJ, 6, e4751. https://doi.org/10.7717/peerj.4751 |
| 22 | Marine mammal | Humpback whale | *Megaptera novaeangliae* | Least Concern | No significative | Pinger | Harcourt, R., Pirotta, V., Heller, G., Peddemors, V., Slip, D. (2014). A whale alarm fails to deter migrating humpback whales: an empirical test. Endangered Species Research, 25(1), 35-42. https://doi.org/10.3354/esr00614 |
| 23 | Marine mammal | Harbour porpoise | *Phocoena phocoena* | Least Concern | NA | Pinger | Culik, B., von Dorrien, C., Müller, V., Conrad, M. (2015). Synthetic communication signals influence wild harbour porpoise (*Phocoena phocoena*) behaviour. Bioacoustics, 24(3), 201-221. https://doi.org/10.1080/09524622.2015.1023848 |
| 24 | Seabird | Guanay cormorant | *Phalacrocorax bougainvillii* | Near Threatened | 85.1% | LED lights | Mangel, J. C., Wang, J., Alfaro-Shigueto, J., Pingo, S., Jimenez, A., Carvalho, F., ... Godley, B. J. (2018). Illuminating gillnets to save seabirds and the potential for multi-taxa bycatch mitigation. Royal Society open science, 5(7), 180254. https://doi.org/10.1098/rsos.180254 |
| | Elasmobranch | Pacific guitarfish | *Rhinobatos planiceps* | Vulnerable | No significative | | |
| | | NA | *Ray spp.* | NA | No significative | | |
| 25 | Marine mammal | Harbour porpoise | *Phocoena phocoena* | Least Concern | 22.2% | Pinger | Larsen, F., Eigaard, O. R. (2014). Acoustic alarms reduce bycatch of harbour porpoises in Danish North Sea gillnet fisheries. Fisheries Research, 153, 108-112. https://doi.org/10.1016/j.fishres.2014.01.010 |
| 26 | Marine mammal | NA | *Delphinus spp.* | NA | 63.6% | Pinger | Mangel, J. C., Alfaro-Shigueto, J., Witt, M. J., Hodgson, D. J., Godley, B. J. (2013). Using pingers to reduce bycatch of small cetaceans in Peru's small-scale driftnet fishery. Oryx, 47(4), 595-606. https://doi.org/10.1017/S0030605312000658 |
| | | Dusky dolphin | *Lagenorhynchus obscurus* | Least Concern | 18.2% | | |
| | | Bottlenose dolphin | *Tursiops truncatus* | Least Concern | 43.8% | | |
| | | NA | *Globicephala spp.* | NA | 100% | | |
| | | Burmesiter porpoise | *Phocoena spinnipinnis* | Near Threatened | 40% | | |
| | Elasmobranch | NA | *Shark spp.* | NA | NA | | |
| | | NA | *Ray spp.* | NA | NA | | |
| 27 | Elasmobranch | Blue shark | *Prionace glauca* | Near Threatened | NA | Rare earth magnet | Porsmoguer, S. B., Bănaru, D., Boudouresque, C. F., Dekeyser, I., Almarcha, C. (2015). Hooks equipped with magnets can increase catches of blue shark (*Prionace glauca*) by longline fishery. Fisheries Research, 172, 345-351. https://doi.org/10.1016/j.fishres.2015.07.016 |

| | | | | | | | |
|---|---|---|---|---|---|---|---|
| 28 | Elasmobranch | Blind shark | *Brachaelurus waddi* | Least Concern | 31% | Ferrite magnet | Richards, R. J., Raoult, V., Powter, D. M., Gaston, T. F. (2018). Permanent magnets reduce bycatch of benthic sharks in an ocean trap fishery. Fisheries Research, 208, 16-21. https://doi.org/10.1016/j.fishres.2018.07.006 |
| 29 | Marine mammal | Bottlenose dolphin | *Tursiops truncatus* | Least Concern | NA | Pinger | Santana-Garcon, J., Wakefield, C. B., Dorman, S. R., Denham, A., Blight, S., Molony, B. W., Newman, S. J. (2018). Risk versus reward: interactions, depredation rates, and bycatch mitigation of dolphins in demersal fish trawls. Canadian Journal of Fisheries and Aquatic Sciences, 75(12), 2233-2240. https://doi.org/10.1139/cjfas-2017-0203 |
| 30 | Sea turtle | Green turtle | *Chelonia mydas* | Endangered | 54% | Physical model (predator) | Wang, J. H., Fisler, S., Swimmer, Y. (2010). Developing visual deterrents to reduce sea turtle bycatch in gill net fisheries. Marine Ecology Progress Series, 408, 241-250. https://doi.org/10.3354/meps08577 |
| | | | | | 40% | Chemical lightsticks | |
| | | | | | 59% | LED lights | |
| 31 | Elasmobranch | Spiny dogfish | *Squalus acanthias* | Vulnerable | 28.2% | SMART hook | O'Connell, C. P., He, P., Joyce, J., Stroud, E. M., Rice, P. H. (2014). Effects of the SMART™ (Selective Magnetic and Repellent-Treated) hook on spiny dogfish catch in a longline experiment in the Gulf of Maine. Ocean & coastal management, 97, 38-43. https://doi.org/10.1016/j.ocecoaman.2012.08.002 |
| | | Blue shark | *Prionace glauca* | Near Threatened | 25% | | |
| | | Porbeagle | *Lamna nasus* | Vulnerable | 100% | | |
| | | Barndoor skate | *Dipturus laevis* | Least Concern | 31.3% | | |
| | | Thorny skate | *Amblyraja radiata* | Vulnerable | 66.7% | | |
| | | Winter skate | *Leucoraja ocellata* | Endangered | 50% | | |
| | | Smooth skate | *Malacoraja senta* | Vulnerable | 100% | | |
| | | Great torpedo ray | *Torpedo nobiliana* | Least Concern | 100% | | |
| 32 | Sea turtle | Green turtle | *Chelonia mydas* | Endangered | 39.7% | UV-LED | Wang, J., Barkan, J., Fisler, S., Godinez-Reyes, C., Swimmer, Y. (2013). Developing ultraviolet illumination of gillnets as a method to reduce sea turtle bycatch. Biology letters, 9(5), 20130383. https://doi.org/10.1098/rsbl.2013.0383 |
| 33 | Seabird | Laysan albatross | *Phoebastria immutabilis* | Near Threatened | No significative | Tori line | Sato, N., Ochi, D., Minami, H., Yokawa, K. (2012). Evaluation of the effectiveness of light streamer tori-lines and characteristics of bait attacks by seabirds in the western North Pacific. PLOS one, 7(5), e37546. https://doi.org/10.1371/journal.pone.0037546 |
| | | Black-footed albatross | *Phoebastria nigripes* | Near Threatened | No significative | | |
| 34 | Elasmobranch | Australian swellshark | *Cephaloscyllium laticeps* | Least Concern | NA | Rare earth magnet | Westlake, E. L., Williams, M., Rawlinson, N. (2018). Behavioural responses of draughtboard sharks (*Cephaloscyllium laticeps*) to rare earth magnets: implications for shark bycatch management within the Tasmanian southern rock lobster fishery. Fisheries Research, 200, 84-92. https://doi.org/10.1016/j.fishres.2018.01.001 |

| | | | | | | | |
|---|---|---|---|---|---|---|---|
| 35 | Sea turtle | Green turtle | *Chelonia mydas* | Endangered | 63.9% | LED lights | Ortiz, N., Mangel, J. C., Wang, J., Alfaro-Shigueto, J., Pingo, S., Jimenez, A., ... Godley, B. J. (2016). Reducing green turtle bycatch in small-scale fisheries using illuminated gillnets: the cost of saving a sea turtle. Marine Ecology Progress Series, 545, 251-259. https://doi.org/10.3354/meps11610 |
| 36 | Seabird | White-chinned petrel | *Procellaria aequinoctialis* | Vulnerable | NA | Mackerel bait replacing squid | González, A., Vega, R., Barbieri, M. Á., Yáñez, E. (2012). Determinación de los factores que inciden en la captura incidental de aves marinas en la flota palangrera pelágica chilena. Latin american journal of aquatic research, 40(SPECISSUE), 786-799. http://dx.doi.org/103856/vol40-issue3-fulltext-25 |
| 37 | Elasmobranch | Blue shark | *Prionace glauca* | Near Threatened | No significative | Electropositive metal alloy | Godin, A. C., Wimmer, T., Wang, J. H., Worm, B. (2013). No effect from rare-earth metal deterrent on shark bycatch in a commercial pelagic longline trial. Fisheries Research, 143, 131-135. https://doi.org/10.1016/j.fishres.2013.01.020 |
| | | Shortfin Mako | *Isurus oxyrinchus* | Endangered | NA | | |
| | | Porbeagle | *Lamna nasus* | Vulnerable | No significative | | |
| 38 | Seabird | Greater shearwater | *Ardenna gravis* | Least Concern | NA | Mackerel bait replacing squid | Li, Y., Browder, J. A., Jiao, Y. (2012). Hook effects on seabird bycatch in the United States Atlantic pelagic longline fishery. Bulletin of Marine Science, 88(3), 559-569.https://doi.org/10.5343/bms.2011.1039 |
| | | Cory's shearwater | *Calonectris borealis* | Least Concern | NA | | |
| | | Northern gannet | *Morus bassanus* | Least Concern | NA | | |
| | | European herring gull | *Larus argentatus* | Least Concern | NA | | |
| 39 | Marine mammal | Humpback whale | *Megaptera novaeangliae* | Least Concern | No significative | Pinger | Pirotta, V., Slip, D., Jonsen, I. D., Peddemors, V. M., Cato, D. H., Ross, G., Harcourt, R. (2016). Migrating humpback whales show no detectable response to whale alarms off Sydney, Australia. Endangered Species Research, 29(3), 201-209. https://doi.org/10.3354/esr00712 |
| 40 | Seabird | Southern Royal albatross | *Diomedea epomophora* | Vulnerable | 100% | Tori line | Domingo, A., Jiménez, S., Abreu, M., Forselledo, R., Yates, O. (2017). Effectiveness of tori line use to reduce seabird bycatch in pelagic longline fishing. PloS one, 12(9), e0184465. https://doi.org/10.1371/journal.pone.0184465 |
| | | Northern Royal albatross | *Diomedea sanfordi* | Endangered | 0% | | |
| | | White-capped albatross | *Thalassarche steadi* | Near Threatened | 100% | | |
| | | Black-browed albatross | *Thalassarche melanophris* | Endangered | 88.5% | | |
| | | Southern giant petrel | *Macronectes giganteus* | LeastConcern | 100% | | |
| | | White-chinned petrel | *Procellaria aequinoctialis* | Vulnerable | 75% | | |
| | | Great shearwater | *Ardenna gravis* | Least Concern | 100% | | |
| 41 | Elasmobranch | Scalloped hammerhead | *Sphyrna lewini* | Critically Endangered | 61.5% | Electropositive metal alloy | Hutchinson, M., Wang, J. H., Swimmer, Y., Holland, K., Kohin, S., Dewar, H., ... Martinez, J. (2012). The effects of a lanthanide metal alloy on shark catch rates. Fisheries Research, 131, 45-51. |
| | | Sandbar shark | *Carcharhinus plumbeus* | Endangered | 18.2% | | |
| | | Tiger shark | *Galeocerdo cuvier* | Near Threatened | 20% | | |
| | | Shortfin Mako | *Isurus oxyrinchus* | Endangered | 0% | | |

| | | | | | | | |
|---|---|---|---|---|---|---|---|
| | | Blue shark | *Prionace glauca* | Near Threatened | 12% | | https://doi.org/10.1016/j.fishres.2012.07.006 |
| | | Pelagic thresher | *Alopias pelagicus* | Endangered | 0% | | |
| | | Blue shark | *Prionace glauca* | Near Threatened | 0% | | |
| 42 | Sea turtle | Olive ridley turtle | *Lepidochelys olivacea* | Vulnerable | 85% | Mackerel bait replacing squid | Santos, M. N., Coelho, R., Fernandez-Carvalho, J., Amorim, S. (2012). Effects of hook and bait on sea turtle catches in an equatorial Atlantic pelagic longline fishery. Bulletin of Marine Science, 88(3), 683-701. https://doi.org/10.5343/bms.2011.1065 |
| | | Leatherback turtle | *Dermochelys coriacea* | Vulnerable | 85% | | |
| | | Loggerhead turtle | *Caretta caretta* | Vulnerable | | | |
| | | Kemp's ridley turtle | *Lepidochelys kempii* | Critically Endangered | | | |
| 43 | Elasmobranch | Blue shark | *Prionace glauca* | Near Threatened | NA | Mackerel bait replacing squid | Coelho, R., Santos, M. N., Amorim, S. (2012). Effects of hook and bait on targeted and bycatch fishes in an equatorial Atlantic pelagic longline fishery. Bulletin of Marine Science, 88(3), 449-467. https://doi.org/10.5343/bms.2011.1064 |
| | | Shortfin Mako | *Isurus oxyrinchus* | Endangered | NA | | |
| | | Smooth hammerhead | *Sphyrnazygaena* | Vulnerable | NA | | |
| | | Silky shark | *Carcharhinus falciformis* | Vulnerable | NA | | |
| | | Bigeye thresher | *Alopias superciliosus* | Vulnerable | NA | | |
| | | Common thresher | *Alopias vulpinus* | Vulnerable | NA | | |
| | | Crocodile shark | *Pseudocarcharias kamoharai* | Least Concern | NA | | |
| | | Longfin Mako | *Isurus paucus* | Endangered | NA | | |
| | | Scalloped hammerhead | *Sphyrna lewini* | Critically Endangered | NA | | |
| | | Oceanic whitetip shark | *Carcharhinus longimanus* | Critically Endangered | NA | | |
| | | Tiger shark | *Galeocerdo cuvier* | Near Threatened | NA | | |
| | | NA | *Myliobatidae spp* | NA | NA | | |
| | | Pelagic stingray | *Pteroplatytrygon violacea* | Least Concern | NA | | |
| 44 | Sea turtle | Leatherback turtle | *Dermochelys coriacea* | Vulnerable | 89% | LED lights | Allman, P., Agyekumhene, A., Stemle, L. (2021). Gillnet illumination as an effective measure to reduce sea turtle bycatch. Conservation Biology, 35(3), 967-975. https://doi.org/10.1111/cobi.13647 |
| | | Olive ridleyturtle | *Lepidochelys olivacea* | Vulnerable | 78% | | |
| | | Green turtle | *Chelonia mydas* | Endangered | 100% | | |
| 45 | Sea turtle | Loggerhead turtle | *Caretta caretta* | Vulnerable | NA | TED | Lucchetti, A., Punzo, E., Virgili, M. (2016). Flexible Turtle Excluder Device (TED): an effective tool for Mediterranean coastal multispecies bottom trawl fisheries. Aquatic Living Resources, 29(2), 201. https://doi.org/10.1051/alr/2016016 |
| 46 | Sea turtle | Leatherback turtle | *Dermochelys coriacea* | Vulnerable | 55% | Two circle hooks | Coelho, R., Santos, M. N., Fernandez-Carvalho, J., Amorim, S. (2015). Effects of hook and bait in a tropical northeast Atlantic pelagic longline fishery: part I—incidental sea turtle bycatch. Fisheries Research, 164, |
| | | Loggerhead turtle | *Caretta caretta* | Vulnerable | 50-59% | | |

| | | | | | | | |
|---|---|---|---|---|---|---|---|
| | | Olive ridley turtle | *Lepidochelys olivacea* | Vulnerable | 50-59% | and mackerel bait | 302-311. https://doi.org/10.1016/j.fishres.2014.11.008 |
| | | Kemp's ridley turtle | *Lepidochelys kempii* | Critically Endangered | 50-59% | | |
| 47 | Sea turtle | Leatherback turtle | *Dermochelys coriacea* | Vulnerable | 65% | Circle hook | Sales, G., Giffoni, B. B., Fiedler, F. N., Azevedo, V. G., Kotas, J. E., Swimmer, Y., Bugoni, L. (2010). Circle hook effectiveness for the mitigation of sea turtle bycatch and capture of target species in a Brazilian pelagic longline fishery. Aquatic Conservation: Marine and Freshwater Ecosystems, 20(4), 428-436. https://doi.org/10.1002/aqc.1106 |
| | | Loggerhead turtle | *Caretta caretta* | Vulnerable | 55% | | |
| 48 | Seabird | Laysan albatross | *Phoebastria immutabilis* | Near Threatened | NA | Tori line (with offal discard and blue-dyed bait) | Gilman, E., Chaloupka, M., Ishizaki, A., Carnes, M., Naholowaa, H., Brady, C., ... Kingma, E. (2021). Tori lines mitigate seabird bycatch in a pelagic longline fishery. Reviews in Fish Biology and Fisheries, 31, 653-666. https://doi.org/10.1007/s11160-021-09659-7 |
| | | Black-footed albatross | *Phoebastria nigripes* | Near Threatened | NA | | |
| 49 | Seabird | Southern Royal albatross | *Diomedea epomophora* | Vulnerable | 100% | BLW | Jiménez, S., Domingo, A., Forselledo, R., Sullivan, B. J., Yates, O. (2019). Mitigating bycatch of threatened seabirds: the effectiveness of branch line weighting in pelagic longline fisheries. Animal Conservation, 22(4), 376-385.https://doi.org/10.1111/acv.12472 |
| | | Black-browed albatross | *Thalassarche melanophris* | Least Concern | 40% | | |
| | | Northern giant petrel | *Macronectes halli* | Least Concern | 0% | | |
| 50 | Seabird | Black-browed albatross | *Thalassarche melanophris* | Least Concern | NA | BLW | Santos, R. C., Silva-Costa, A., Sant'Ana, R., Gianuca, D., Yates, O., Marques, C., Neves, T. (2019). Improved line weighting reduces seabird bycatch without affecting fish catch in the Brazilian pelagic longline fishery. Aquatic Conservation: Marine and Freshwater Ecosystems, 29(3), 442-449. https://doi.org/10.1002/aqc.3002 |
| | | White-chinned petrel | *Procellaria aequinoctialis* | Vulnerable | NA | | |
| | | Great shearwater | *Ardenna gravis* | Least Concern | NA | | |
| 51 | Seabird | Cape petrel | *Daption capense* | Least Concern | NA | Tori line (with BLW) | Sato, N., Katsumata, N., Yokota, K., Uehara, T., Fusejima, I., Minami, H. (2016). Tori-lines with weighted branch lines reduce seabird bycatch in eastern South Pacific longline fishery. Aquatic Conservation: Marine and Freshwater Ecosystems, 26(1), 95-107. https://doi.org/10.1002/aqc.2492 |
| | | White-chinned petrel | *Procellaria aequinoctialis* | Vulnerable | NA | | |
| | | Westland petrel | *Procellaria westlandica* | Endangered | NA | | |
| 52 | Seabird | NA | *Diomedea spp.* | NA | NA | Tori line | Yokota, K., Minami, H., Kiyota, M. (2011). Effectiveness of tori-lines for further reduction of incidental catch of seabirds in pelagic longline fisheries. Fisheries Science, 77, 479-485. https://doi.org/10.1007/s12562-011-0357-4 |
| | | NA | *Thalassarche spp.* | NA | NA | | |
| | | NA | *Phoebetria spp.* | NA | NA | | |
| | | NA | *Macronectes spp.* | NA | NA | | |
| | | NA | *Procellaria spp.* | NA | NA | | |
| | | NA | *Puffinus spp.* | NA | NA | | |
| 53 | Seabird | Laysan albatross | *Phoebastria inmutabilis* | Near Threatened | NA | Tori line | Sato, N., Minami, H., Katsumata, N., Ochi, D., Yokawa, K. (2013). Comparison of the effectiveness of paired and single tori lines for preventing bait attacks by seabirds and their bycatch in pelagic longline fisheries. Fisheries research, 140, 1419.https://doi.org/10.1016/j.fishres.2012.11.007 |
| | | NA | *Larus spp.* | NA | NA | | |

| | | | | | | | |
|---|---|---|---|---|---|---|---|
| 54 | Seabird | Northern gannet | *Morus bassanus* | Least Concern | NA | High contrast panels<br>Black hooks<br>Scarybird | Oliveira, N., Almeida, A., Alonso, H., Constantino, E., Ferreira, A., Gutierrez, I., ... Andrade, J. (2021). A contribution to reducing bycatch in a high priority area for seabird conservation in Portugal. Bird Conservation International, 31(4), 553-572. https://doi.org/10.1017/S0959270920000489 |
| | | Cory's shearwater | *Calonectris borealis* | Least Concern | NA | | |
| | | European shag | *Gulosus aristotelis* | Least Concern | NA | | |
| | | Great cormorant | *Phalacrocorax carbo* | Least Concern | NA | | |
| | | Great shearwater | *Ardenna gravis* | Least Concern | NA | | |
| | | Yellow-legged gull | *Larus michahellis* | Least Concern | NA | | |
| | | Lesser black-backed gull | *Larus fuscus* | Least Concern | NA | | |
| 55 | Seabird | White-chinned petrel | *Procellaria aequinoctialis* | Vulnerable | 97.7% | Tori line | Paterson, J. R., Yates, O., Holtzhausen, H., Reid, T., Shimooshili, K., Yates, S., ... Wanless, R. M. (2019). Seabird mortality in the Namibian demersal longline fishery and recommendations for best practice mitigation measures. Oryx, 53(2), 300-309. https://doi.org/10.1017/S0030605317000230 |
| | | Atlantic yellow-nosed albatross | *Thalassarche chlororhynchos* | Endangered | 100% | | |
| | | Brown skua | *Catharacta antarctica* | Least Concern | 100% | | |
| | | Black-browed albatross | *Thalassarche melanophris* | Least Concern | 100% | | |
| | | Sooty shearwater | *Ardenna grisea* | Near Threatened | 100% | | |
| | | Cape gannet | *Morus capensis* | Endangered | 100% | | |
| 56 | Marine mammal | Harbour porpoise | *Phocoena phocoena* | Least Concern | NA | Pinger | Van Beest, F. M., Kindt-Larsen, L., Bastardie, F., Bartolino, V., Nabe-Nielsen, J. (2017). Predicting the population-level impact of mitigating harbor porpoise bycatch with pingers and time-area fishing closures. Ecosphere, 8(4), e01785. https://doi.org/10.1002/ecs2.1785 |
| 57 | Seabird | Scopoli's shearwater | *Calonectris diomedea* | Least Concern | 83.3% | Night setting | Cortes, V., Gonzalez-Solis, J. (2018). Seabird bycatch mitigation trials in artisanal demersal longliners of the Western Mediterranean. PloS one, 13(5), e0196731.https://doi.org/10.1371/journal.pone.0196731 |
| | | Balearic shearwater | *Puffinus mauretanicus* | Critically Endangered | 100% | | |
| | | Yelkouan shearwater | *Puffinus yelkouan* | Vulnerable | 100% | | |
| | | Yellow-legged gull | *Larus michahellis* | Least Concern | 100% | | |
| | | Audouin's gull | *Larus audouinii* | Vulnerable | 100% | | |
| 58 | Seabird | White-chinned petrel | *Procellaria aequinoctialis* | Vulnerable | NA | Night setting<br>BLW<br>Tori line<br>Thawed bait<br>Bird exclusion device<br>Offal discard | Collins, M. A., Hollyman, P. R., Clark, J., Söffker, M., Yates, O., Phillips, R. A. (2021). Mitigating the impact of longline fisheries on seabirds: Lessons learned from the South Georgia Patagonian toothfish fishery (CCAMLR Subarea 48.3). Marine Policy, 131, 104618. https://doi.org/10.1016/j.marpol.2021.104618 |
| | | Black-browed albatross | *Thalassarche melanophris* | Least Concern | NA | | |
| | | Grey-headed albatross | *Thalassarche chrysostoma* | Endangered | NA | | |
| | | NA | *Macronectes spp.* | NA | NA | | |
| | | Bigeye thresher | *Alopias superciliosus* | Vulnerable | 0% | | |
| | | Common thresher | *Alopias vulpinus* | Vulnerable | | | |
| | | Porbeagle | *Lamna nasus* | Vulnerable | 41.2% | | |
| | | Copper shark | *Carcharhinus brachyurus* | Near Threatened | 87.5% | | |

| | | | | | | | |
|---|---|---|---|---|---|---|---|
| 59 | Elasmobranch | Dusky shark | *Carcharhinus obscurus* | Vulnerable | 0% | Tori line and BLW | Jiménez, S., Forselledo, R., Domingo, A. (2019). Effects of best practices to reduce seabird bycatch in pelagic longline fisheries on other threatened, protected and bycaught megafauna species. Biodiversity and Conservation, 28(13), 3657-3667. https://doi.org/10.1007/s10531-019-01842-4 |
| | | Night shark | *Carcharhinus signatus* | Endangered | 89.5% | | |
| | | Smooth hammerhead | *Sphyrna zygaena* | Vulnerable | 86.3% | | |
| | | Scalloped hammerhead | *Sphyrna lewini* | Endangered | 100% | | |
| | | Pelagic stingray | *Pteroplatytrygon violacea* | Least Concern | 27.8% | | |
| | | Spinetail devil ray | *Mobula mobular* | Endangered | 50% | | |
| | | Bentfin devil ray | *Mobula thurstoni* | Endangered | | | |
| | Sea turtle | Loggerhead turtle | *Caretta caretta* | Vulnerable | 18.1% | | |
| | | Leatherback turtle | *Dermochelys coriacea* | Vulnerable | 73.3% | | |
| | Marine mammal | NA | *Arctocephalus spp.* | NA | NA | | |
| 60 | Elasmobranch | Scalloped hammerhead | *Sphyrna lewini* | Endangered | NA | TED | Gupta, T., Booth, H., Arlidge, W., Rao, C., Manoharakrishnan, M., Namboothri, N., ... Milner-Gulland, E. J. (2020). Mitigation of elasmobranch bycatch in trawlers: a case study in Indian fisheries. Frontiers in Marine Science, 7, 571. https://doi.org/10.3389/fmars.2020.00571 |
| | | Sharpnose guitarfish | *Glaucostegus granulatus* | Critically Endangered | NA | | |
| | | Widenose guitarfish | *Glaucostegus obtusus* | Critically Endangered | NA | | |
| 61 | Marine mammal | Australian snubfin dolphin | *Orcaella heinsohni* | Vulnerable | NA | Pinger | Soto, A. B., Cagnazzi, D., Everingham, Y., Parra, G. J., Noad, M., Marsh, H. (2013). Acoustic alarms elicit only subtle responses in the behaviour of tropical coastal dolphins in Queensland, Australia. Endangered Species Research, 20(3), 271-282.https://doi.org/10.3354/esr00495 |
| | | Indo-Pacific humpback dolphin | *Sousa chinensis* | Vulnerable | NA | | |
| 62 | Seabird | Shy albatross | *Thalassarche cauta* | Near Threatened | 20.5% | Night setting<br>Tori line<br>BLW<br>Thawed bait | Rollinson, D. P., Wanless, R. M., Ryan, P. G. (2017). Patterns and trends in seabird bycatch in the pelagic longline fishery off South Africa. African Journal of Marine Science, 39(1), 9-25. https://doi.org/10.2989/1814232X.2017.1303396 |
| | | Black-browed albatross | *Thalassarche melanophris* | Least Concern | 6.8% | | |
| | | Indian yellow-nosed albatross | *Thalassarche carteri* | Endangered | 3.3% | | |
| | | Atlantic yellow-nosed albatross | *Thalassarche chlororhynchos* | Endangered | 0.8% | | |
| | | Northern Royal albatross | *Diomedea sanfordi* | Endangered | 0.1% | | |
| | | Wandering albatross | *Diomedea exulans* | Vulnerable | 0.2% | | |
| | | Northern giant petrel | *Macronectes halli* | Least Concern | 0.3% | | |
| | | White-chinned petrel | *Procellaria aequinoctialis* | Vulnerable | 65.7% | | |
| | | Grey petrel | *Procellaria cinerea* | Near Threatened | 0.1% | | |
| | | Cape petrel | *Daption capense* | Least Concern | 0.1% | | |

| | | | | | | | |
|---|---|---|---|---|---|---|---|
| | | Great shearwater | *Ardenna gravis* | Least Concern | 0.1% | | |
| | | Brown skua | *Catharacta antarctica* | Least Concern | 0.1% | | |
| | | Cape gannet | *Morus capensis* | Endangered | 1.9% | | |
| | | King penguin | *Aptenodytes patagonicus* | Least Concern | 0.1% | | |
| 63 | Elasmobranch | Cuckoo ray | *Leucoraja naevus* | Least Concern | NA | Ticker | Kynoch, R. J., Fryer, R. J., Neat, F. C. (2015). A simple technical measure to reduce bycatch and discard of skates and sharks in mixed-species bottom-trawl fisheries. ICES Journal of Marine Science, 72(6), 1861-1868. https://doi.org/10.1093/icesjms/fsv037 |
| | | Flapper skate | *Dipturus intermedius* | Critically Endangered | NA | | |
| | | Spotted ray | *Raja montagui* | Least Concern | NA | | |
| | | Thornback ray | *Raja clavata* | Near Threatened | NA | | |
| | | Blackmouth dogfish | *Galeus melastomus* | Least Concern | NA | | |
| | | Lesser-spotted dogfish | *Scylorhinus canicula* | Least Concern | 2.3% | | |
| | | Spiny dogfish | *Squalus acanthias* | Vulnerable | NA | | |
| 64 | Marine mammal | Bottlenose dolphin | *Tursiops truncatus* | Least Concern | NA | Bycatch excluder device | Jaiteh, V. F., Allen, S. J., Meeuwig, J. J., Loneragan, N. R. (2013). Subsurface behavior of bottlenose dolphins (*Tursiops truncatus*) interacting with fish trawl nets in northwestern Australia: Implications for bycatch mitigation. Marine Mammal Science, 29(3), E266-E281. https://doi.org/10.1111/j.1748-7692.2012.00620.x |
| 65 | Sea turtle | Loggerhead turtle | *Caretta caretta* | Vulnerable | 68% | Buoyless net | Peckham, S. H., Lucero-Romero, J., Maldonado-Díaz, D., Rodríguez-Sánchez, A., Senko, J., Wojakowski, M., Gaos, A. (2016). Buoyless nets reduce sea turtle bycatch in coastal net fisheries. Conservation Letters, 9(2), 114-121. https://doi.org/10.1111/conl.12176 |
| | | Green turtle | *Chelonia mydas* | Endangered | | | |
| | | Olive ridley turtle | *Lepidochelys olivacea* | Vulnerable | | | |
| 66 | Marine mammal | Harbour porpoise | *Phocoena phocoena relicta* | Endangered | NA | Pinger | Popov, D. V., Meshkova, G. D., Hristova, P. D., Gradev, G. Z., Rusev, D. Z., Panayotova, M. D., Dimitrov, H. A. (2020). Pingers as Cetacean Bycatch Mitigation Measure in Bulgarian Turbot Fishery. Acta Zoologica Bulgarica, Supplement, 15, 235-242. |
| | | Bottlenose dolphin | *Tursiops truncatus ponticus* | Endangered | NA | | |
| 67 | Marine mammal | Harbour porpoise | *Phocoena phocoena* | Least Concern | 96.9% | Pinger | Moan, A. N. D. R. É., Bjørge, A. R. N. E. (2021). Pinger trials in Norwegian commercial fisheries confirm that pingers reduce harbour porpoise bycatch rates and demonstrate low level of pinger-associated negative impacts on day-to-day fishing operations. IWC Scientific Committee. Report number: SC, 68. |
| | | Harbor seal | *Phoca vitulina* | Least Concern | 33.5% | | |
| 68 | Sea turtle | Green turtle | *Chelonia mydas* | Endangered | 48.8% | LED lights | Kakai, T. M. (2019). Assessing the effectiveness of LED lights for the reduction of sea turtle bycatch in an artisanal gillnet fishery-a case study from the north coast of Kenya. Western Indian Ocean Journal of Marine Science, 18(2), 37-44. https://doi.org/10.4314/wiojms.v18i2.4 |
| | | Hawksbill turtle | *Eretmochelys imbricata* | Critically Endangered | 44.4% | | |
| | | Loggerhead turtle | *Caretta caretta* | Vulnerable | 20% | | |
| | | Olive ridley turtle | *Lepidochelys olivacea* | Vulnerable | 100% | | |

| | | | | | | | |
|---|---|---|---|---|---|---|---|
| 69 | Seabird | Black-browed albatross | *Thalassarche melanophris* | Least Concern | NA | BLW | Sullivan, B. J., Kibel, B., Kibel, P., Yates, O., Potts, J. M., Ingham, B., ... Wanless, R. M. (2018). At-sea trialling of the Hookpod: a 'one-stop'mitigation solution for seabird bycatch in pelagic longline fisheries. Animal Conservation, 21(2), 159-167. https://doi.org/10.1111/acv.12388 |
| | | White-chinned petrel | *Procellaria equinoctialis* | Vulnerable | NA | | |
| | | Southern Royal albatross | *Diomedea epomophora* | Vulnerable | NA | | |
| | | Shy albatross | *Thalassarche cauta* | Near Threatened | NA | | |
| 70 | Sea turtle | Loggerhead turtle | *Caretta caretta* | Vulnerable | NA | TED | Sala, A., Lucchetti, A., Affronte, M. (2011). Effects of Turtle Excluder Devices on bycatch and discard reduction in the demersal fisheries of Mediterranean Sea. Aquatic Living Resources, 24(2), 183-192. https://doi.org/10.1051/alr/2011109 |
| 71 | Sea turtle | Loggerhead turtle | *Caretta caretta* | Vulnerable | 97% | TED | Warden, M. L. (2011). Modeling loggerhead sea turtle (*Caretta caretta*) interactions with US Mid-Atlantic bottom trawl gear for fish and scallops, 2005–2008. Biological Conservation, 144(9), 2202-2212.https://doi.org/10.1016/j.biocon.2011.05.012 |
| 72 | Sea turtle | Olive ridley turtle | *Lepidochelys olivacea* | Vulnerable | 100% | TED | Raghu Prakash, R., Boopendranath, M.R., Vinod, M. (2016). Performance evaluation of Turtle Excluder Device off Dharma in Bay of Bengal, Fish. Technol., 53(3): 183-189. |
| 73 | Sea turtle | Leatherback turtle | *Dermochelys coriacea* | Vulnerable | 100% | TED | Casale, P., Abitsi, G., Aboro, M. P., Agamboue, P. D., Agbode, L., Allela, N. L., ... Formia, A. (2017). A first estimate of sea turtle bycatch in the industrial trawling fishery of Gabon. Biodiversity and conservation, 26, 2421-2433. https://doi.org/10.1007/s10531-017-1367-z |
| | | Olive ridley turtle | *Lepidochelys olivacea* | Vulnerable | 100% | | |
| | | Hawksbill turtle | *Eretmochelys imbricata* | Critically Endangered | 100% | | |
| | | Green turtle | *Chelonia mydas* | Endangered | 100% | | |

### 2.2. Species and areas

The species of sea turtles included in this review are classified by the UICN Red List of Threatened Species (UICN, 2023) as: Vulnerable (VU), such as the loggerhead turtle (*Caretta caretta*) (Casale and Tucker, 2017), Endangered (EN), such as the green turtle (*Chelonia mydas*) (Seminoff, 2004), and Critically endangered (CR), such as the hawksbill turtle (*Eretmochelys imbricata*) (Mortimer and Donnelly, 2008). It is worth noting that only one species of sea turtle, the flatback turtle (*Natator depressus*) (Standards and Subcommittee, 1996), is classified as Data deficient (DD). One of the main threats to this species is fisheries, especially gillnet and longline fisheries, which are predominant in American countries such as Mexico, Peru, Ecuador or Brazil.

Marine mammals are also included in the review, as is the case of odontocetes and mysticetes, Sirenian and Pinniped. Several species of marine mammals have been classified by the UICN Red List of Threatened Species (UICN, 2023). These include the harbour porpoise (*Phocoena phocoena*) as LC (Braulik et al. 2020), burmeister's porpoise (*Phocoena spinipinnis*) as NT (Félix et al. 2018), the dugong (*Dugong dugon*) as VU (Marsh and Sobtzick, 2019) and the Black Sea bottlenose dolphin (*Tursiops truncatus ponticus*) (Birkun, 2012) and harbour porpoise subpopulation (*Phocoena phocoena relicta*) (Birkun and Frantzis, 2008) as EN. The predominant type of fishery with which they interact is the gillnet fishery of European countries such as Denmark, Sweden, Germany and Norway, among others.

The seabird species included are classified by the UICN Red List of Threatened Species (UICN, 2023) as LC, NT, VU, EN and CR, with the waved albatross (*Phoebastria irrorata*) (International, 2018) and Balearic shearwater (BirdLife International, 2018a) being highly endangered. With regard to seabird species, the fishery with the highest amount of by-catch is the longline fishery, in which the birds become hooked on the baited hooks. In this case, the countries in which studies have been carried out in an attempt to minimize the by-catch of these species are more widely distributed throughout the world, specifically in five of the six continents (Asia, Africa, America, Europe and Oceania).

Finally, the species of elasmobranchs have been classified by the UICN Red List of Threatened Species (UICN, 2023) as LC, NT, VU, EN and CR, with the largetooth sawfish (*Pristis pristis*) (Espinosa et al. 2022), scalloped hammerhead (*Sphyrna lewini*) (Rigby, Dulvy, et al., 2019), oceanic whitetip shark (*Carcharhinus longimanus*) (Rigby et al. 2019), sharpnose guitarfish (*Glaucostegus granulatus*) (Kyne et al. 2022), widenose guitarfish (*Glaucostegus obtusus*) (Kyne and Jabado, 2021) and flapper skate (*Dipturus intermedius*) (Ellis et al. 2021) being highly threatened. The fishery with the highest by-catch for this group is the longline fishery, and the country in which most by-catch mitigation measures have been studied is Australia, where the main type of fishing is pot/trap.

**Table 2 SI.** Conservation status of some of the threatened species from each group (sea turtles, marine mammals, seabirds and elasmobranchs).

| ENDANGERED SPECIES | | | | | |
|---|---|---|---|---|---|
| **GROUP** | **COMMON NAME** | **SCIENTIFIC NAME** | **STATUS CONSERVATION UICN** | **PHOTO** | **REFERENCE** |
| **SEA TURTLES** | Loggerhead turtle | *Caretta caretta* | Vulnerable | 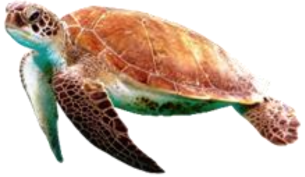 | Casale, P., Tucker, A. D. (2017). *Caretta caretta*. The IUCN Red List of Threatened Species. https://dx.doi.org/10.2305/IUCN.UK.2017-2.RLTS.T3897A119333622.en |
| | Green turtle | *Chelonia mydas* | Endangered | 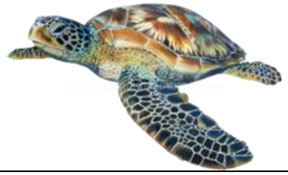 | Seminoff, J. A. (2004). *Chelonia mydas*. The IUCN Red List of Threatened Species. https://dx.doi.org/10.2305/IUCN.UK.2004.RLTS.T4615A11037468.en |
| | Hawksbill turtle | *Eretmochelys imbricata* | Critically Endangered | 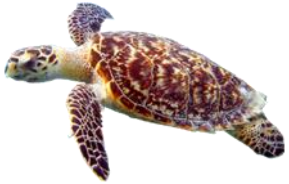 | Mortimer, J. A., Donnelly, M. (2008). *Eretmochelys imbricata*. The IUCN Red List of Threatened Species. https://dx.doi.org/10.2305/IUCN.UK.2008.RLTS.T8005A12881238.en |
| | Flatback turtle | *Natator depressus* | Data Deficient | 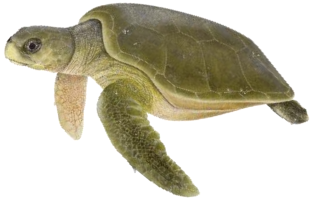 | Standards, R. L., Subcommittee, P. (1996). *Natator depressus*. The IUCN Red List of Threatened Species.<br><br>Photo: Dawn Witherington<br>SWOT (2009). Discovering the flatback Australia`s own sea turtle. The State of the World's Sea Turtles. Volume IV, p.1-52. https://static1.squarespace.com/static/5b80290bee1759a50e3a86b3/t/5bb3f7e10852290ffa4791fa/1538521077015/swot4.pdf |

| | | | | | |
|---|---|---|---|---|---|
| **MARINE MAMMALS** | Harbour porpoise | *Phocoena phocoena* | Least Concern | 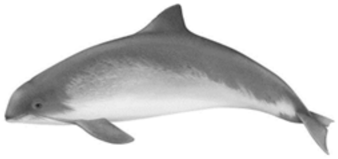 | Braulik, G., Minton, G., Amano, M., Bjørge, A. (2020). *Phocoena phocoena*. The IUCN Red List of Threatened Species. https://dx.doi.org/10.2305/IUCN.UK.2020-2.RLTS.T17027A50369903.en<br><br>Photo: Lucy Molleson<br>Carlén, I., Nunny, L., Simmonds, M. P. (2021). Out of sight, out of mind: how conservation is failing European porpoises. Frontiers in Marine Science, 8, 13. https://doi.org/10.3389/fmars.2021.617478 |
| | Burmeister's porpoise | *Phocoena spinipinnis* | Near Threatened | 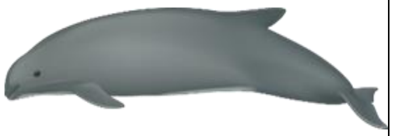 | Félix, F., Alfaro, J., Reyes, J., Mangel, J., Dellabianca, N., Heinrich, S., Crespo, E. (2018). *Phocoena spinipinnis*. The IUCN Red List of Threatened Species. https://dx.doi.org/10.2305/IUCN.UK.2018-2.RLTS.T17029A50370481.en |
| | Dugong | *Dugong dugon* | Vulnerable | 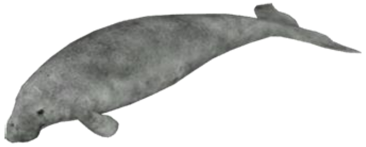 | Marsh, H., Sobtzick, S. (2019). *Dugong dugon*. The IUCN Red List of Threatened Species. https://dx.doi.org/10.2305/IUCN.UK.2015-4.RLTS.T6909A160756767.en |
| | Black bottlenose dolphin subpopulation | *Tursiops truncatus ponticus* | Endangered | 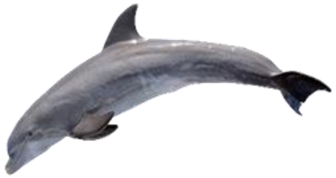 | Birkun, A. (2012). *Tursiops truncatus ponticus*. The IUCN Red List of Threatened Species. https://dx.doi.org /10.2305/IUCN.UK.2012.RLTS.T133714A17771698.en |
| | Black Sea harbour porpoise | *Phocoena phocoena relicta* | Endangered | 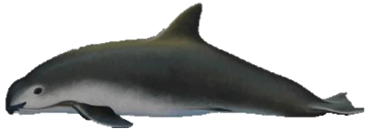 | Birkun Jr., A., Frantzis, A. (2008). *Phocoena phocoena ssp. relicta*. The IUCN Red List of Threatened Species. https://dx.doi.org/10.2305/IUCN.UK.2008.RLTS.T17030A6737111.en. |

| | | | | | |
|---|---|---|---|---|---|
| **SEABIRDS** | Waved albatross | *Phoebastria irrorata* | Critically Endangered | 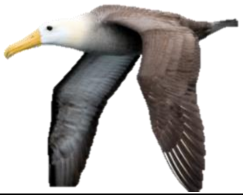 | International, B. (2018a). *Phoebastria irrorata*. The IUCN Red List of Threatened Species. https://dx.doi.org/10.2305/IUCN.UK.2018-2.RLTS.T22698320A132641638.en<br><br>(*) Photo: Brian Sullivan<br>Brian Sullivan (12 de julio de 2017). Waved Albatross (*Phoebastria irrorata*). The CornellLab of Ornithology Macaulay Library. https://ebird.org/species/wavalb?siteLanguage=es |
| | Balearic shearwater | *Puffinus mauretanicus* | Critically Endangered | 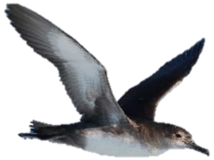 | International, B. (2018b). *Puffinus mauretanicus*. The IUCN Red List of Threatened Species. https://dx.doi.org/10.2305/IUCN.UK.2018-2.RLTS.T22728432A132658315.en<br><br>(*) Photo: Jorge López Álvarez<br>Jorge López (2 de junio de 2019). Pardela balear (*Puffinus mauretanicus*). The CornellLab of Ornithology Macaulay Library. https://ebird.org/species/balshe1?siteLanguage=es_MX |
| **ELASMOBRANCHS** | Largetooth sawfish | *Pristis pristis* | Critically Endangered | 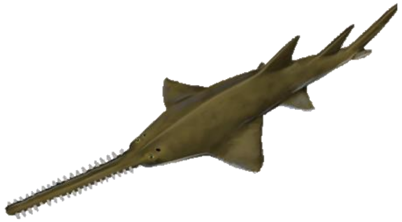 | Espinosa, M., Bonfil-Sanders, R., Carlson, J., Charvet, P., Chevis, M., Dulvy, N. K., Everett, B., Faria, V., Ferretti, F., Fordham, S., Grant, M. I., Haque, A. B., Harry, A. V., Jabado, R. W., Jones, G. C. A., Kelez, S., Lear, K. O., Morgan, D. L., Phillips, N. M., Wueringer, B. E. (2022). *Pristis pristis.* The IUCN Red List of Threatened Species. https://dx.doi.org/10.2305/IUCN.UK.2022-2.RLTS.T18584848A58336780.en<br><br>Photo: NOAA fisheries<br>NOAA fisheries (2023). Largetooth Sawfish. National Oceanic and Atmospheric Administration. https://www.fisheries.noaa.gov/species/largetooth-sawfish/overview |
| | Scalloped hammerhead | *Sphyrna lewini* | Critically Endangered | 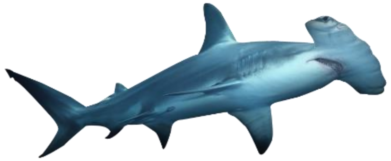 | Rigby, C. L., Dulvy, N. K., Barreto, R., Carlson, J., Fernando, D., Fordham, S., Francis, M. P., Herman, K., Jabado, R. W., Liu, K. M., Marshall, A., Pacoureau, N., Romanov, E., Sherley, R. B., Winker, H. (2019). *Sphyrna lewini*. The IUCN Red List of Threatened Species. |
| | Oceanic whitetip shark | *Carcharhinus longimanus* | Critically Endangered | 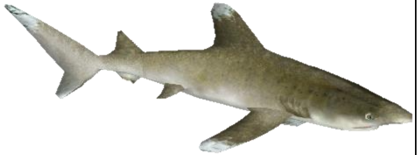 | Rigby, C. L., Barreto, R., Carlson, J., Fernando, D., Fordham, S., Francis, M. P., Herman, K., Jabado, R. W., Liu, K. M., Marshall, A., Pacoureau, N., Romanov, E., Sherley, R. B., Winker, H. (2019). *Carcharhinus longimanus*. The IUCN Red List of Threatened Species. https://dx.doi.org/10.2305/IUCN.UK.2019-3.RLTS.T39374A2911619.en |

| | | | | | |
|---|---|---|---|---|---|
| | Sharpnose guitarfish | *Glaucostegus granulatus* | Critically Endangered | 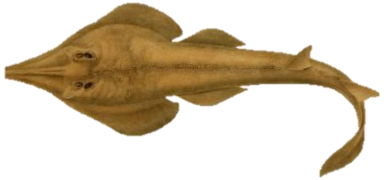 | Kyne, P. M., Haque, A. B., Charles, R., Jabado, R. W. (2022). *Glaucostegus granulatus*. The IUCN Red List of Threatened Species.<br><br>Photo: Johannes Müller and Jacob Henle<br>Müller, J., Henle, J. (1841). Systematische Beschreibung der Plagiostomen. Museum of Comparative Zoology--Biodiversity Heritage Library digitization project. Berlin, Veit und comp, 1841. https://doi.org/10.5962/bhl.title.6906 |
| | Widenose guitarfish | *Glaucostegus obtusus* | Critically Endangered | 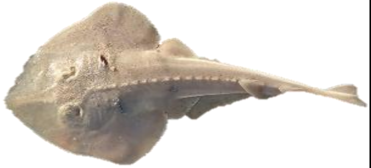 | Kyne, P. M., Jabado, R. W. (2021). *Glaucostegus obtusus*. The IUCN Red List of Threatened Species. https://dx.doi.org/10.2305/IUCN.UK.2021-3.RLTS.T60170A207283191.en<br><br>(*) Photo: Parth Heblekar<br>Parth Heblekar (abril de 2019). *Glaucostegus obtusus*. iNaturalist. https://www.inaturalist.org/observations/22691609 |
| | Flapper skate | *Dipturus intermedius* | Critically Endangered | 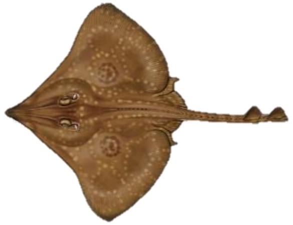 | Ellis, J. R., McCully-Philipps, S. R., Sims, D., Walls, R. H. L., Cheok, J., Derrick, D., Dulvy, N. K. (2021). Dipturus intermedius. The IUCN Red List of Threatened Species. https://dx.doi.org/10.2305/IUCN.UK.2021-2.RLTS.T18903491A68783461.en<br><br>Photo: Mark Dando<br>Mark Dando. (NA). Flapper skate. Orkney Skate Trust. https://www.orkneyskatetrust.co.uk/flapper-skate/ |

Note 1: Unreferenced photographs are free to use as "png" images.

Note 2: The photographs marked with (*) have undergone modifications to the original photograph (removal of the background) in order to homogenize the Table 2SI.

### 3. Results and discussion

#### 3.1. Overview of published literature

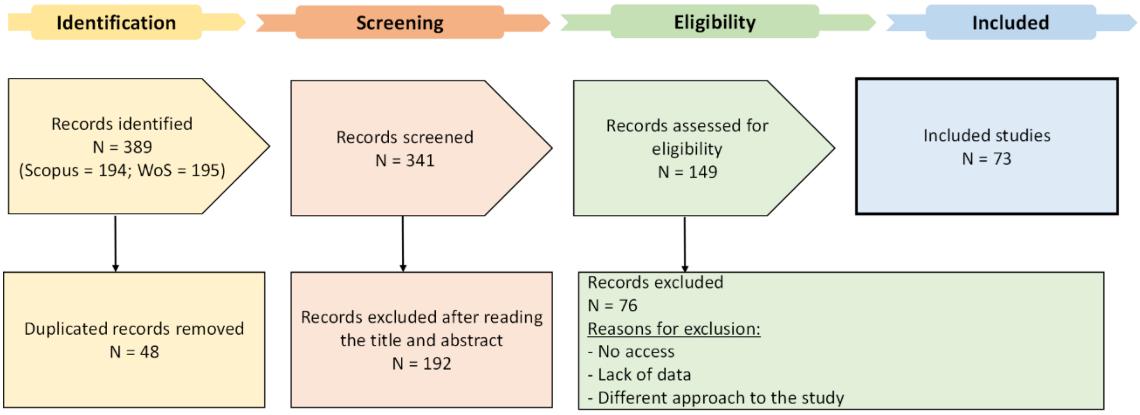


**Figure 1 S1**. Prism diagram showing the different phases and specific exclusion criteria of the papers reviewed for this study. Exclusion criteria: Duplicated records removed during identification phase, records excluded after reading the title and abstract during screening phase and records excluded owing to no access, lack of data and different approach to the study during the eligibility phase.

**Table 3 SI.** Count of the 73 papers found during the literature screening process from 2010 to 2022 and included in the review. The percentage is the proportion of studies published out of all 73 studies found.

| Year | Number of studies | % |
|---|---|---|
| 2010 | 2 | 2.7 |
| 2011 | 4 | 5.5 |
| 2012 | 7 | 9.6 |
| 2013 | 7 | 9.6 |
| 2014 | 4 | 5.5 |
| 2015 | 4 | 5.5 |
| 2016 | 5 | 6.8 |
| 2017 | 6 | 8.2 |
| 2018 | 9 | 12.3 |
| 2019 | 6 | 8.2 |
| 2020 | 9 | 12.3 |
| 2021 | 7 | 9.6 |
| 2022 | 3 | 4.1 |
| TOTAL | 73 | 100 |

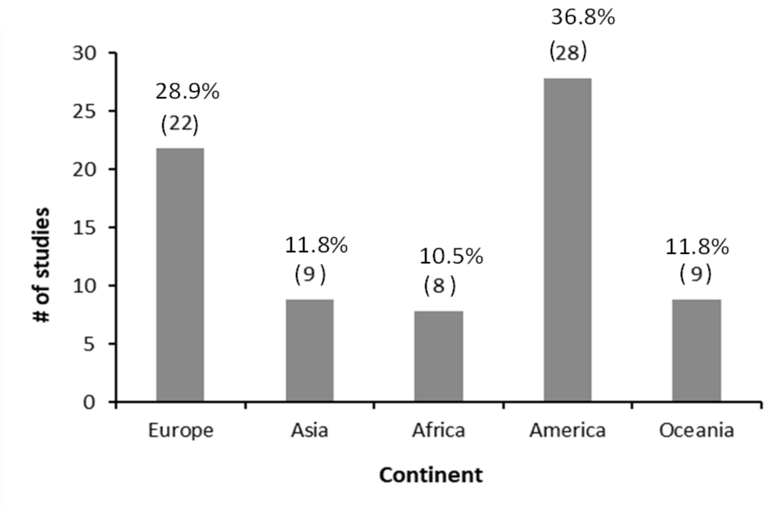


**Figure 2 SI.** Count of studies per continent belonging to the fishing fleet involved in each of the 73 field studies found during the literature screening process. The numbers above the bars are the studies for that continent (A) and the percentage is the proportion of studies published out of all 73 studies found. Note: the sum of the columns is 76 because two papers cover several countries.

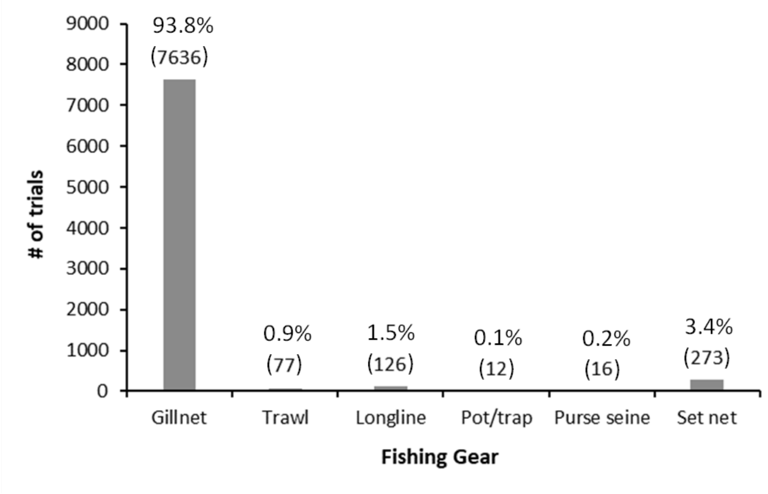


**Figure 3 SI.** Trial count by gear for the total number of 8140 trials in the 31 studies selected after carrying out the literature screening process, since they were those that indicated the number of trials performed. Proofs of concept and laboratory studies have been excluded from this review. Numbers on top of the bars are the trials for each type of fishing gear (A) and the percentage is the proportion of trials published out of all 8140 trials found.

**3.2. Worldwide mitigation measures**

Table 1. Methods SI
